## Supplementary material for "Altered tRNA processing is linked to a distinct and unusual La protein in *Tetrahymena thermophila*": Description of Supplementary Data

### Description of Additional Supplementary Data

**Supplementary Data 1.** TGIRT-sequencing normalized tRNA counts from *Tetrahymena thermophila* samples.

Next generation sequencing tRNA counts are shown as transcripts per million (TPM). Biological triplicates were analysed. MLP1_IPx: size-excluded MLP1-associated RNAs following ribonucleoprotein-immunoprecipitation, WTx: size-excluded total RNA from wild type (WT) strain, MLP1_KOx: size-excluded total RNA from the partial MLP1 knockout (KO) strain. MLP1_WT and MLP1_IP are paired samples obtained from the same biological replicate. Relatively lowly expressed transcripts are shown in grey (TPM < 1).

**Supplementary Data 2.** Input sequences for custom Python script to obtain raw counts for 3'-U ending and 3'-CCA ending tRNAs from raw .fastq files.

*Tetrahymena thermophila* tRNA sequences were screened for unique "fishing" sequences in the 3'-end of the mature tRNA for each isoacceptor (see Summary Table, right) in order to obtain counts for mature tRNAs (ending in -CCA) and pre-tRNAs (ending in different length of -U) (see Table S2 and S3).

**Supplementary Data 3.** Sequence information for pre-tRNA and mature tRNAs from different eukaryotic species.

The genomic sequences *for Homo sapiens, Mus musculus, Drosophila melanogaster, Arabidopsis thaliana, Saccharomyces cerevisiae* and Schizosaccharomyces *pombe* were obtained from the Genomic tRNA Database (GtRNAdb) and *T. thermophila* from the UCSC Genome Browser.
