## Supplementary Data for "Altered tRNA processing is linked to a distinct and unusual La protein in *Tetrahymena thermophila*"

### Supplementary Information for "Altered tRNA processing is linked to a distinct and unusual La protein in *Tetrahymena thermophila*"

Kerkhofs, Kyra[1]; Garg, Jyoti[1]; Fafard-Couture, Étienne[2]; Abou Elela, Sherif[3]; Scott, Michelle[2]; Pearlman, Ronald E.[1]; Bayfield, Mark A.[1*]

[1] Department of Biology, Faculty of Science, York University, Toronto, Ontario M3J 1P3, Canada,

[2] Département de Biochimie et de Génomique Fonctionnelle, Faculté de Médecine et des Sciences de la Santé, Université de Sherbrooke, Sherbrooke, Québec, J1E 4 K8, Canada,

[3] Département de Microbiologie et d'Infectiologie, Faculté de Médecine et des Sciences de la Santé, Université de Sherbrooke, Sherbrooke, Québec, J1E 4 K8, Canada.

**Figure S1.** Multiple sequence alignments of La proteins from several eukaryotes demonstrate high conservation of the LaM and absence of the RRM1.

**Figure S2.** Mlp1 demonstrates preferential binding of certain tRNA isotypes and unprocessed pre-tRNAs.

**Figure S3.** Mlp1 binding to typical La protein target RNAs.

**Figure S4.** Mlp1 binding to uridylate RNA does not discriminate between the position of the uridylate.

**Figure S5.** tRNA mediated suppression protein expression in *Schizosaccharomyces pombe*.

**Figure S6.** Confirmation of the partial Mlp1 *Tetrahymena thermophila* knockout strain.

**Figure S7.** The effect of short 3'-trailer sequences on 5'-leader composition and Mlp1 binding affinity.

**Figure S8.** Primary sequence alignments of the 3'-exonuclease Rex1, 3'-endonuclease RNase Z and LSm2-8 complex from different eukaryotic species.

**Table S1**. Kd values from EMSAs determining binding of Mlp1 to different processed pre-tRNA intermediates and mature tRNA.

**Table S2.** Raw counts for 3'-U ending and 3'-CCA ending tRNAs from raw .fastq files from wild type (WT) input tRNA and Mlp1-immunoprecipitated tRNA (IP).

**Table S3.** Raw counts for 3'-U ending and 3'-CCA ending tRNAs from raw .fastq files from wild type (WT) and partial Mlp1 knockout (KO) strains.

**Table S4.** Number of tRNA genes encoded in the genome of different eukaryotic species for each isotype (top) and isoacceptor (bottom).

**Table S5.** List of oligonucleotides used in this study.

**Table S6**. NCBI accession numbers of primary amino acid sequences used for conservation analysis.

# S1

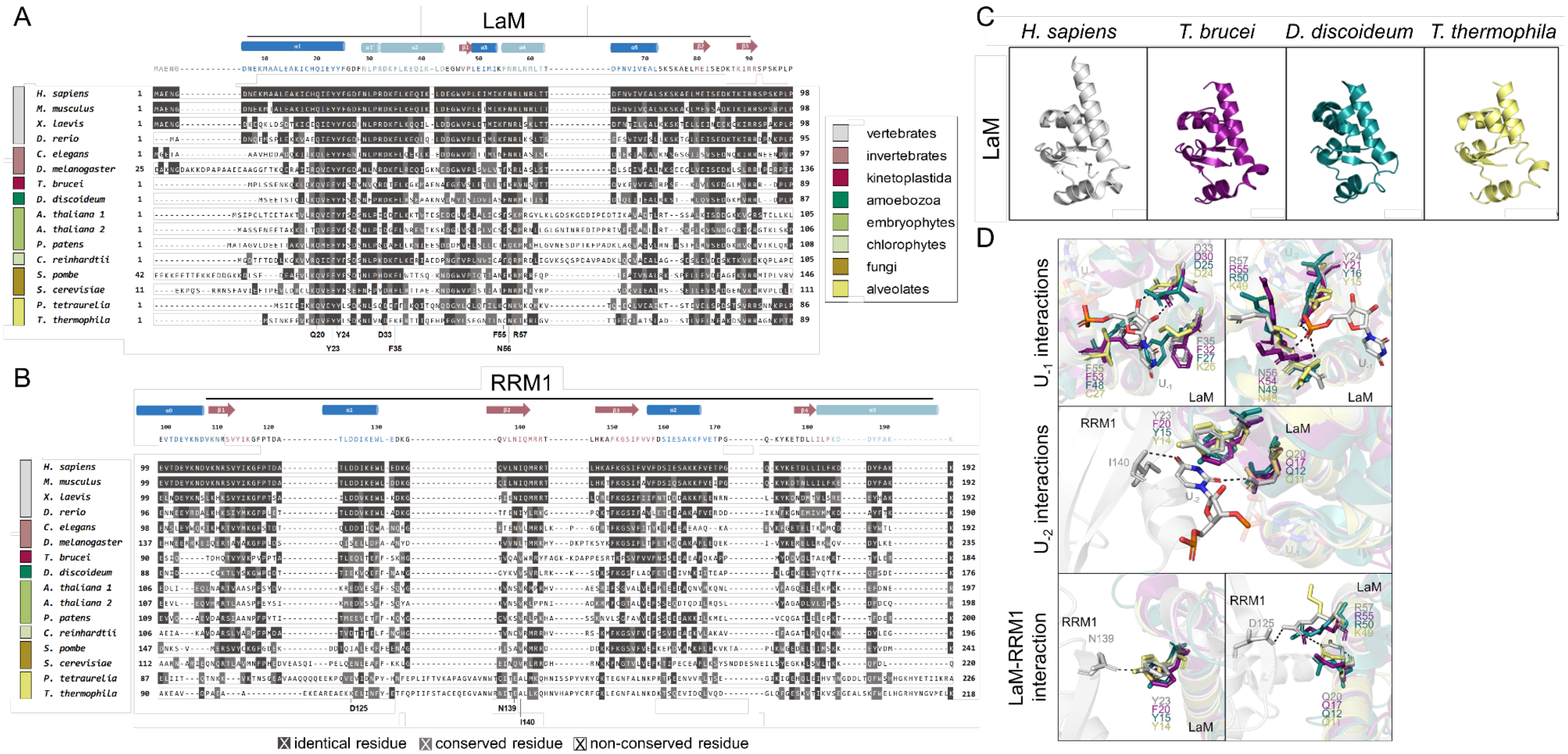


**Figure S1. Multiple sequence alignments of La proteins from several eukaryotes demonstrate high conservation of the La Motif (LaM) and absence of the RNA recognition motif-1 (RRM1).**

(A,B) Primary amino acid alignments of full RNA-binding domains LaM and RRM1 from different eukaryotic lineages. A dark grey background indicates identical residues, light grey conserved residues and white indicates no conservation. Most uridylate binding residues are conserved in the LaM of alveolates, whereas residues in the RRM1 are more variable. Important uridylate binding residues are shown at the bottom. The secondary structure motif of the high-resolution hLa protein structure is shown at the top with β-sheets shown in red and α-helices in blue (dark blue: typical α-helices found in the winged-helix fold and classic RRM, light blue: inserted α-helices found in La proteins specifically) (PDB: 2VOD).

(C) High-resolution structures of the LaM in different species: *Homo sapiens* in white (PDB: 2VOD)[14], *Trypanosoma brucei* in purple (PDB: 1S29)[10], *Dictyostelium discoideum* in teal (PDB: 2M5W)[56] and the predicted tertiary structure *for Tetrahymena thermophila* in yellow[42].

(D) Magnified views of La-$U_{-1}$ RNA, La-$U_{-2}$ and LaM-RRM interactions. $U_{-1}$ is the most 3'-terminal uridylate possessing 2'-OH and 3'-OH ends. Most uridylate binding residues located in the LaM are conserved in *Tetrahymena thermophila,* except for F35 and F55 (hLa numbering) which bind the most 3'-terminal $U_{-1}$. Carbon atoms within the RNA ($U_{-1}$ and $U_{-2}$) from the *Homo sapiens* La structure are colored in white with other atoms colored by type: oxygen in red; nitrogen in blue and phosphorous in orange. Hydrogen bonds are shown as black dashed lines.

# S2

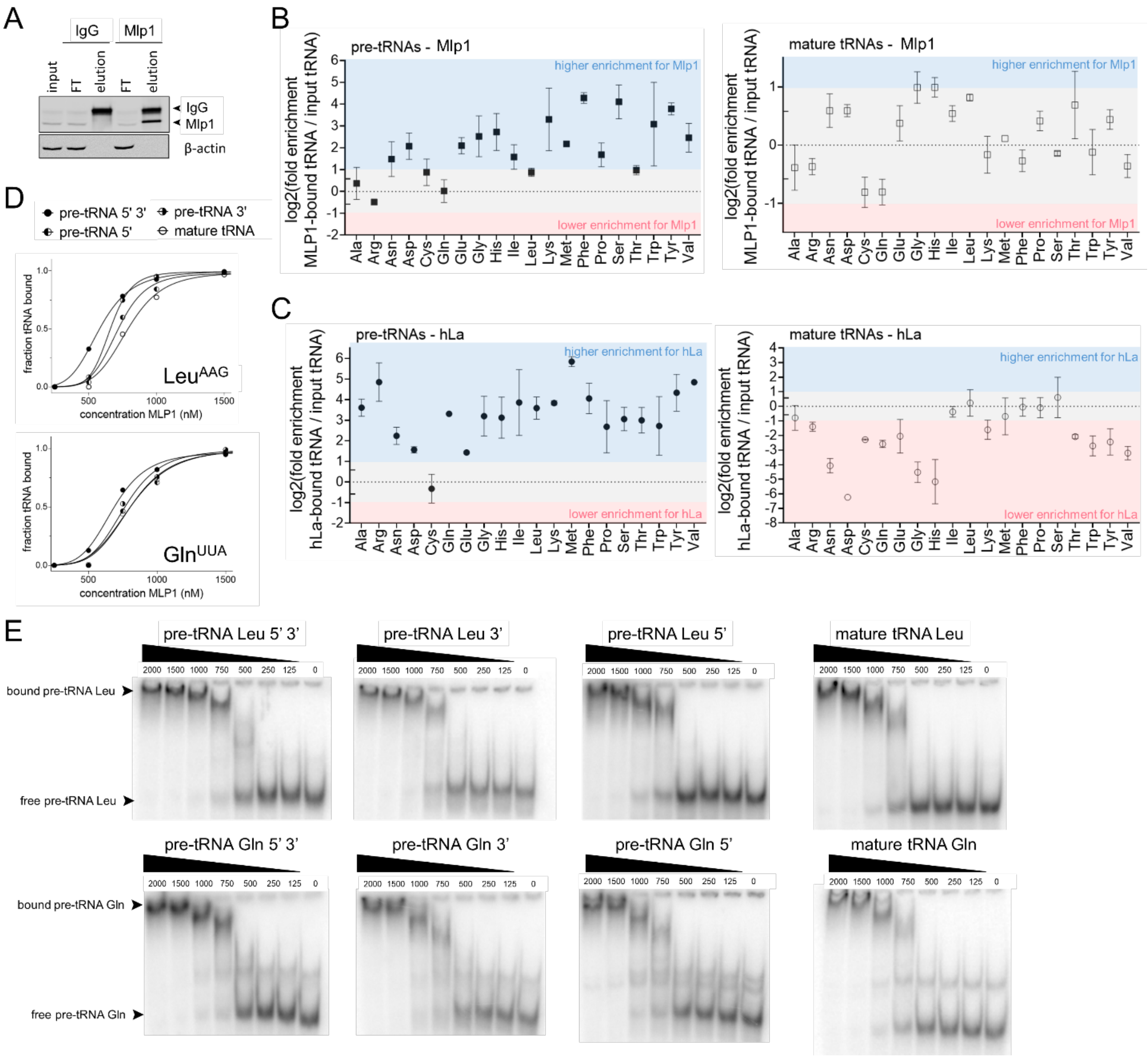


**Figure S2. Mlp1 demonstrates preferential binding of certain tRNA isotypes and unprocessed pre-tRNAs.**
**(A)** Western blot confirming Mlp1-specific immunoprecipitation from *Tetrahymena thermophila* using an affinity purified rabbit anti-Mlp1 antibody and a rabbit isotype immunoglobin G (IgG) control. The Mlp1-specific antibody was used for both ribonucleoprotein-immunoprecipitation (RNP-IP) and western blotting. The heavy chain (50 kDa) of the antibodies was detected with the secondary anti-rabbit antibody and shown as IgG. Loading control: β-actin.
**(B-C)** Next generation sequencing data of tRNAs split by tRNA isotypes for Mlp1 (n=3) (B) and hLa from Gogakos *et al.* (n=2) (C). Fold enrichment is shown as the log2 transformed ratio between Mlp1- or hLa-immunoprecipitated tRNAs and input tRNAs. Both Mlp1 and hLa have a higher binding affinity for pre-tRNAs compared to mature tRNAs.
**(D-E)** Binding curves from EMSAs (D) and corresponding native gels (E) comparing binding affinity of Mlp1 between $^{32}$P-labeled pre-tRNA containing 5'- and 3'-extensions or 5'- or 3'-extensions and mature tRNA for Leu$^{AAG}$ and Gln$^{UUA}$. The highest binding affinity is found for 5'- and 3'-end containing pre-tRNAs. See Table S1 for Kd quantifications.

## S3

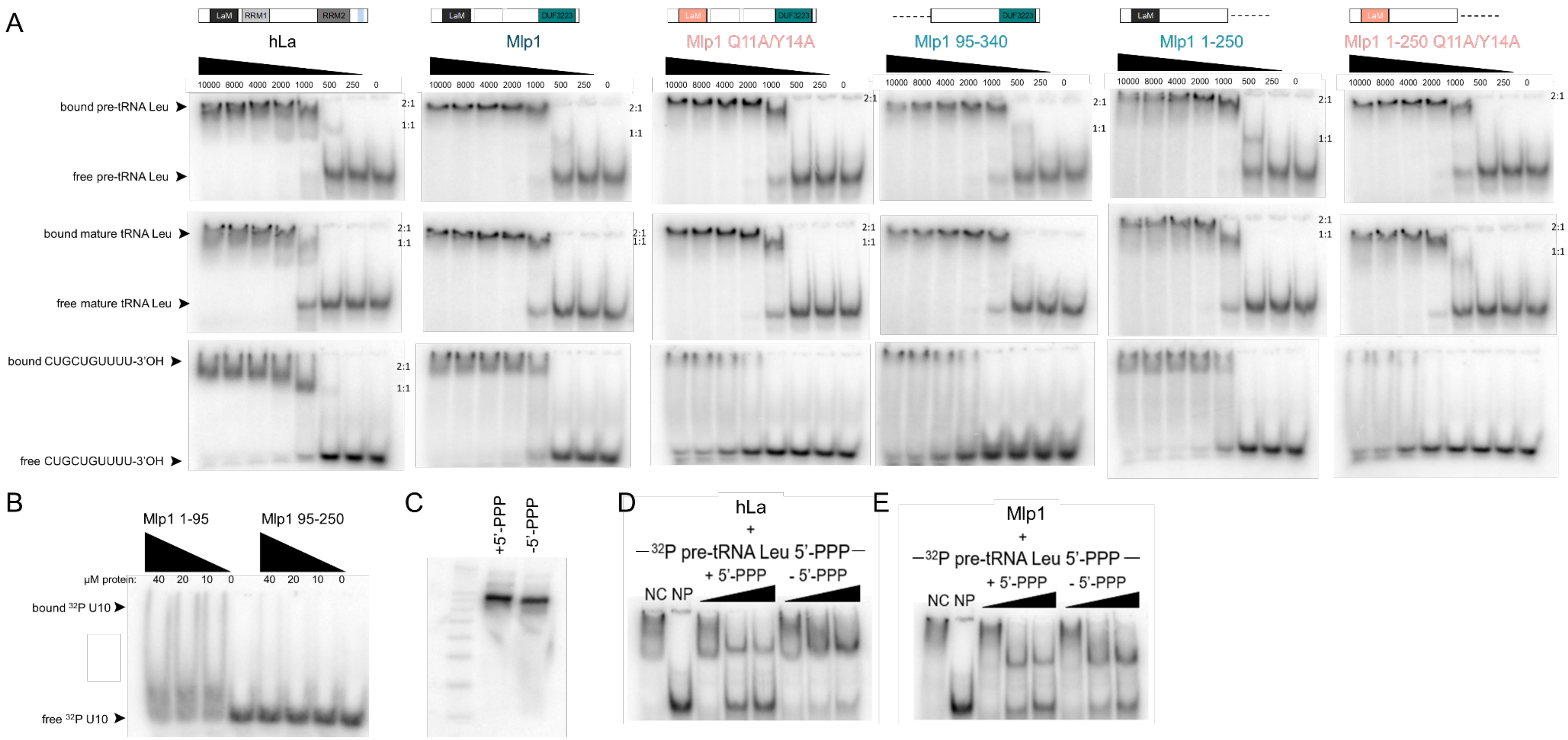


**Figure S3. Mlp1 binding to typical La protein target RNAs.**
**(A)** Native gels comparing binding to $^{32}$P-labeled pre-tRNAs, mature tRNAs and a 3'-trailer sequence CUGCUGUUUU-3'OH. Unbound RNA is found at the bottom of the gel, while protein-RNA complexes shift upwards. The numbering on the side of the gels indicate binding events. A single protein bound to the RNA is indicated at 1:1 and two proteins bound to the same RNA is denoted at 2:1.
**(B)** Native gels comparing binding to $^{32}$P-labeled U10 RNA for mutants Mlp1 1-95 (LaM only) and Mlp1 95-250 (middle domain only).
**(C)** Denaturing gel of *in vitro* transcribed $^{32}$P-labeled 5'-leader containing, 3'-trailer processed pre-tRNAs produced side-by-side as unlabelled-competitor pre-tRNAs (lacking $^{32}$P-labeled labeling) used in (E,F) to confirm successful removal of the 5'-triphosphate group.
**(D,E)** Native gels comparing unlabelled-competitors 5'-triphosphate containing pre-tRNA (+5'PPP) and dephosphorylated pre-tRNA (-5'PPP) for $^{32}$P-labeled +5'PPP binding on hLa (D) or Mlp1 (E). Competition is seen as a decrease in protein-RNA complex and increase in unbound RNA. NP: no protein lane, NC: no unlabeled competitor lane

# S4

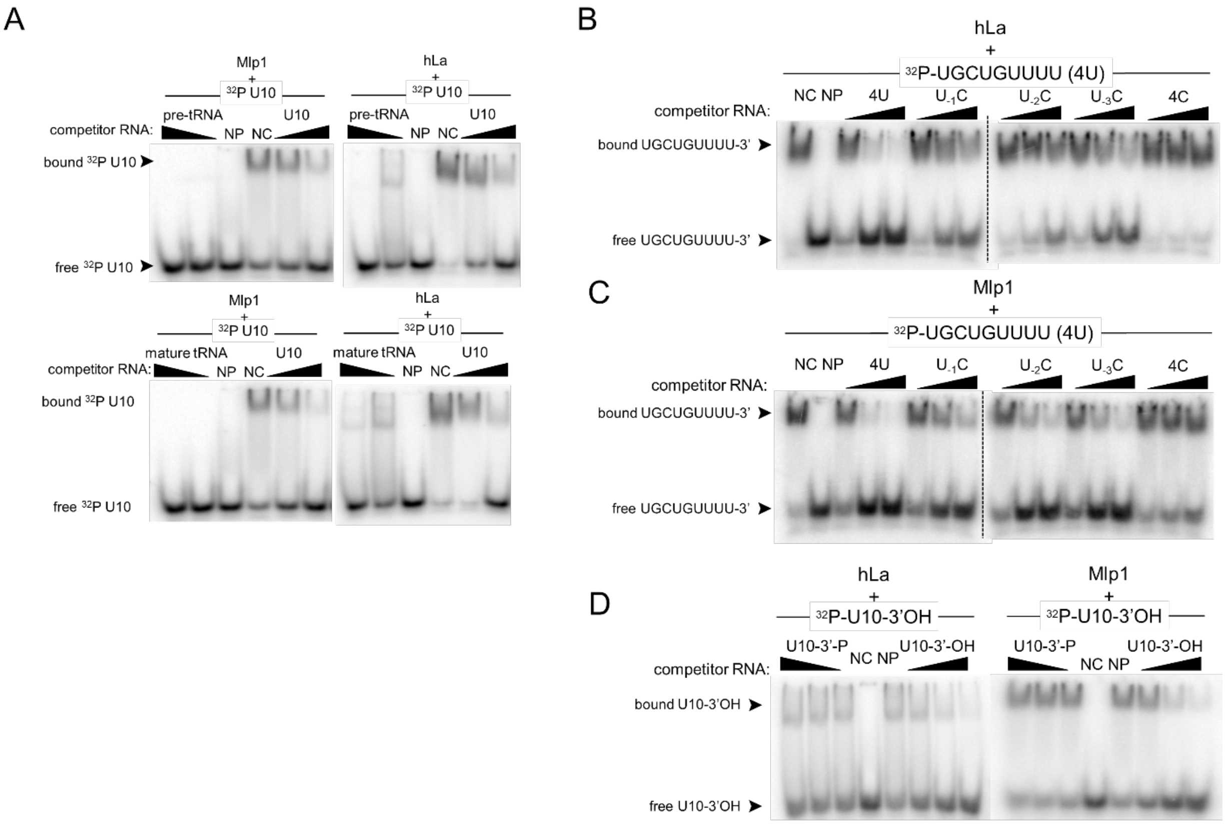


**Figure S4. Mlp1 binding to uridylate RNA does not discriminate between the position of the uridylate.**

**(A)** Native gels comparing unlabelled-competitors U10 (positive control), pre-tRNA and mature tRNA for $^{32}$P-labeled U10 binding. Competition is seen as a decrease in protein-RNA complex and increase in unbound RNA. NP: no protein lane, NC: no competitor lane.

**(B-C)** Native gels comparing unlabelled-competitors CUGCUGUUUU (4U), CUGCUGUUUC ($U_{-1}C$), CUGCUGUUCU ($U_{-2}C$), CUGCUGUCUU ($U_{-3}C$) and CUGCUGCCCC (4C) for $^{32}$P-labeled 4U binding on hLa (B) or Mlp1 (C). Competition is seen as a decrease in protein-RNA complex and increase in unbound RNA. NP: no protein lane, NC: no competitor lane.

**(D)** Native gels comparing a 3'-phosphorylated substrate (U10-3'-P) and normal 3'-OH containing substrate (U10-3'OH) for binding of $^{32}$P-labeled U10-3'-OH. Competition is seen as a decrease in protein-RNA complex and increase in unbound RNA. No competition was observed with the phosphorylated target for either Mlp1 or hLa. NP: no protein lane, NC: no competitor lane.

# S5

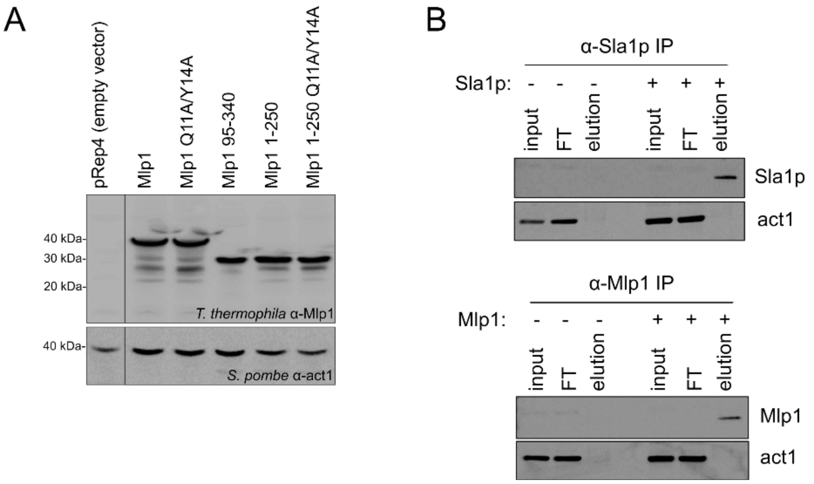


**Figure S5. tRNA mediated suppression protein expression in *Schizosaccharomyces pombe*.**
**(A)** Western blot confirming Mlp1 protein expression levels in *Schizosaccharomyces pombe* following pRep4 plasmid transformation in the tRNA-mediated suppression assay Figure 4A. Loading control: act1.
**(B)** Western blot confirming immunoprecipitation (IP) of Sla1p and Mlp1 from Sla1p- and Mlp1-transformed *Schizosaccharomyces pombe* ySH9 tRNA suppressor strains. pRep4 empty vector transformed strains were used as a control for background immunoprecipitation using Sla1p and Mlp1 antibodies. Loading control: act1. FT: flow through.

# S6

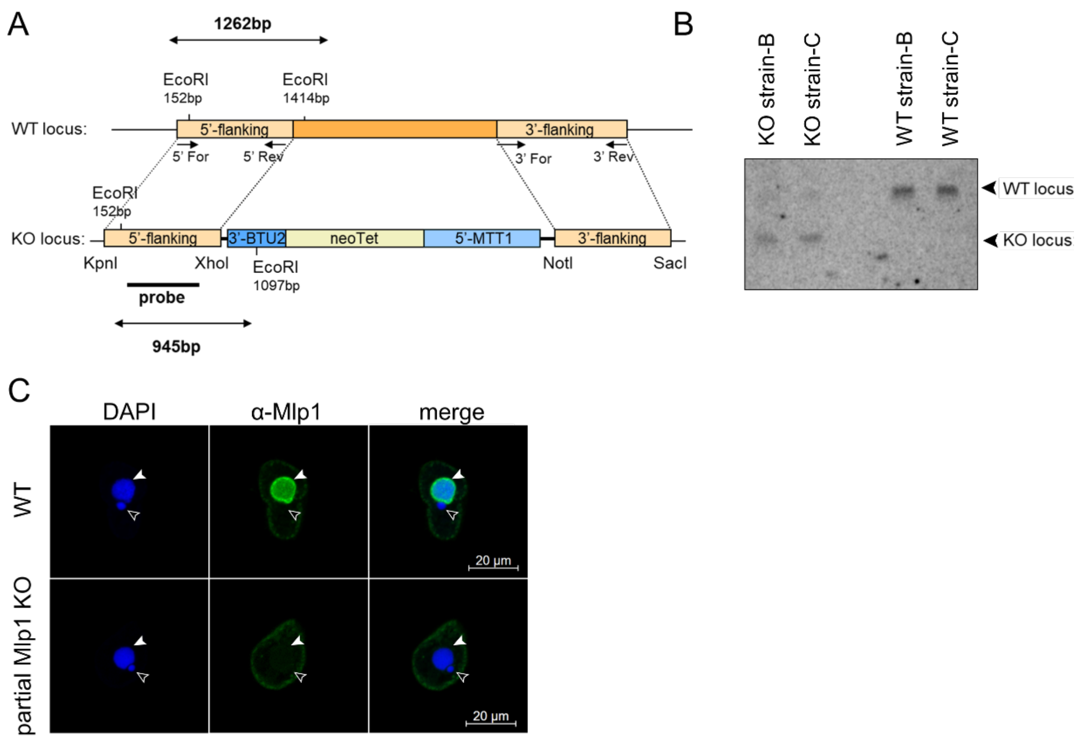


**Figure S6. Confirmation of the partial Mlp1 *Tetrahymena thermophila* knockout strain.**
**(A)** Schematic overview of the *Tetrahymena thermophila* wild type (WT) locus encoding endogenous Mlp1 and knockout (KO) locus following homologous recombination between 5'- and 3'-flanking regions of the genome with transformation plasmid encoding the neomycin (neoTet) selection marker. Restriction enzymes used for cloning 5'- and 3'-flanking regions for homologous recombination are shown below the KO locus. Genomic DNA digest before southern blotting were performed with *EcoRI* restriction enzyme generating a 1262 bp fragment for the WT locus and a 945 bp fragment for the KO locus.
**(B)** Southern blot of genomic DNA digested with restriction enzyme *EcoRI* probed with $^{32}$P-labeled PCR-generated probe reveals almost complete KO of Mlp1 (see schematic in A).
**(C)** Indirect immunofluorescent staining of *Tetrahymena thermophila* WT and partial Mlp1 KO strains using the polyclonal rabbit anti-Mlp1 antibody. Nuclei were stained using DAPI. Full white arrows are denoting the transcriptionally active macronucleus and empty white arrows show the transcriptionally inactive micronucleus.

# S7

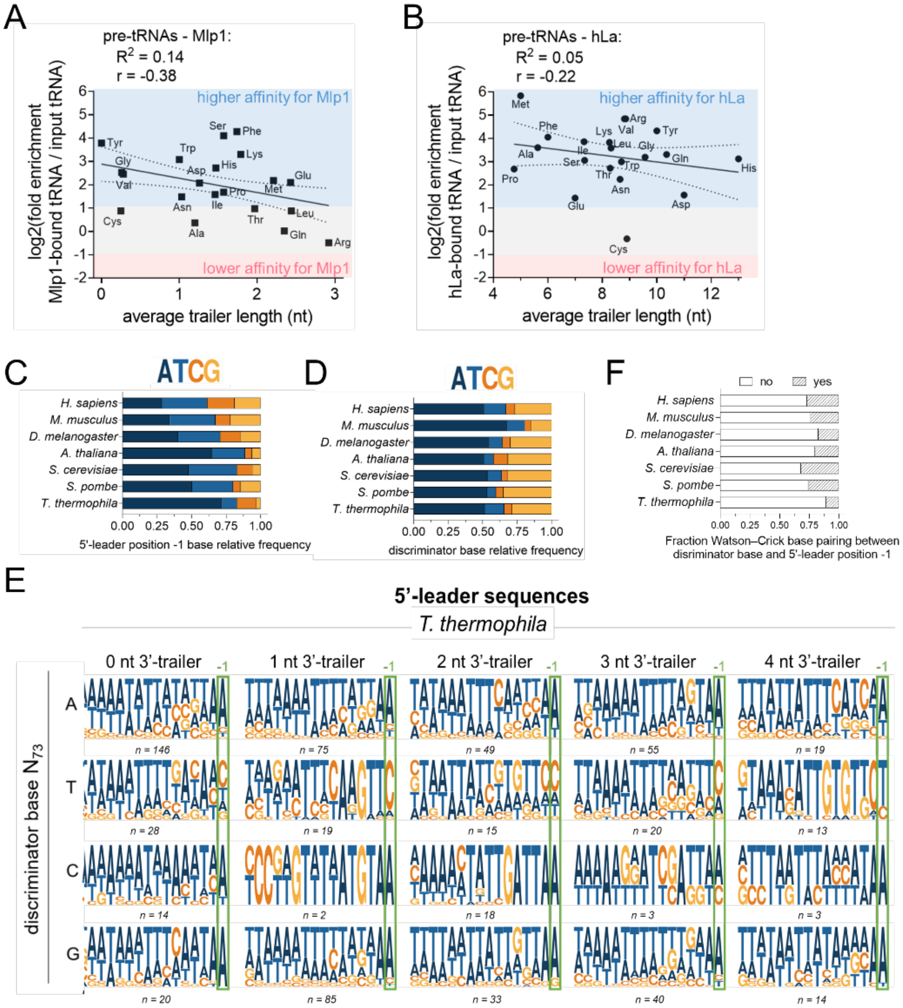


**Figure S7. The effect of short 3'-trailer sequences on 5'-leader composition and Mlp1 binding affinity.**
**(A,B)** Next generation sequencing data of Mlp1-bound pre-tRNAs (n=3) (A) and hLa-bound pre-tRNAs (n=2) (B) split by tRNA isotypes compared against 3'-trailer lengths.
**(C)** Distribution of the nucleotide at the most 3'-terminal position in the 5'-leader of pre-tRNAs from different eukaryotes. *Tetrahymena thermophila* has a high frequency of adenosines in this position.
**(D)** Distribution of the nucleotide found as the discriminator base ($N_{73}$) in different eukaryotes. Distribution is equal between different species.
**(E)** Logo analysis of *Tetrahymena thermophila* 5'-leader sequences split by the discriminator base identity and 3'-trailer length. The number of analyzed pre-tRNAs is shown underneath each logo.
**(F)** Distribution of pre-tRNAs containing a perfect Watson-Crick base pairing between the discriminator base ($N_{73}$) preceding the 3'-trailer and most 3'-terminal 5'-leader nucleotide ($N_{-1}$).

# S8

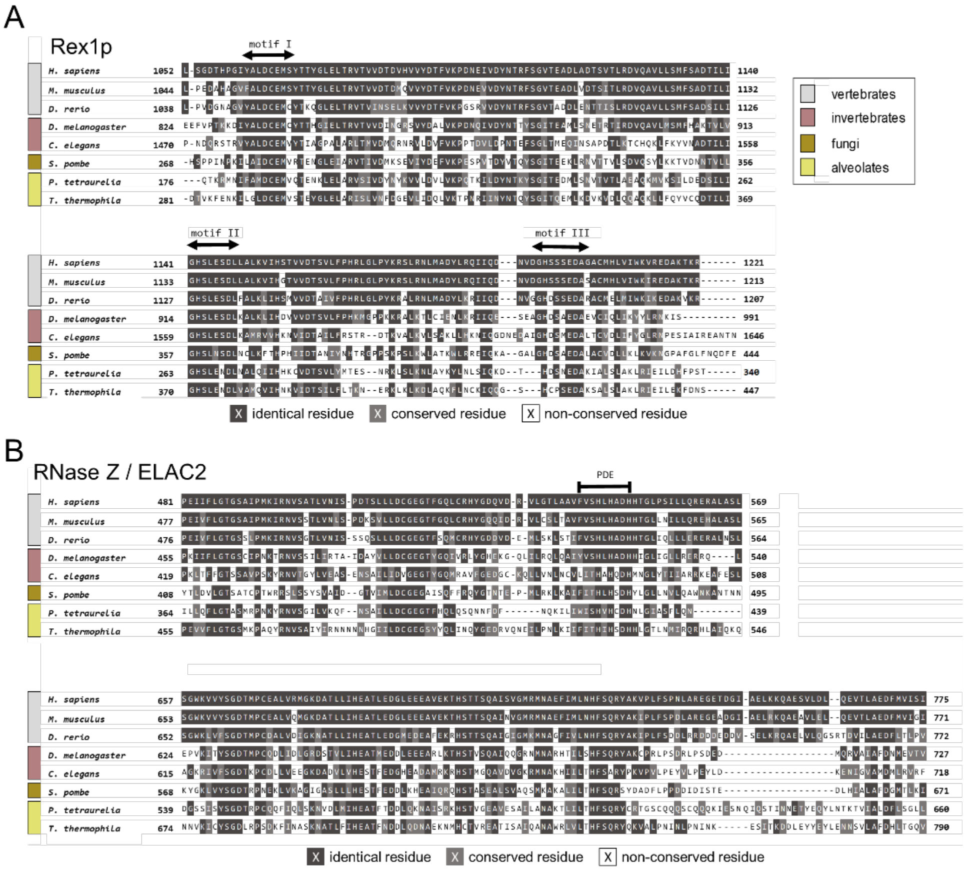
A
Rex1p
motif I
motif II
motif III
H. sapiens
M. musculus
D. rerio
D. melanogaster
C. elegans
S. pombe
P. tetraurelia
T. thermophila
vertebrates
invertebrates
fungi
alveolates
identical residue
conserved residue
non-conserved residue
B
RNase Z / ELAC2
PDE

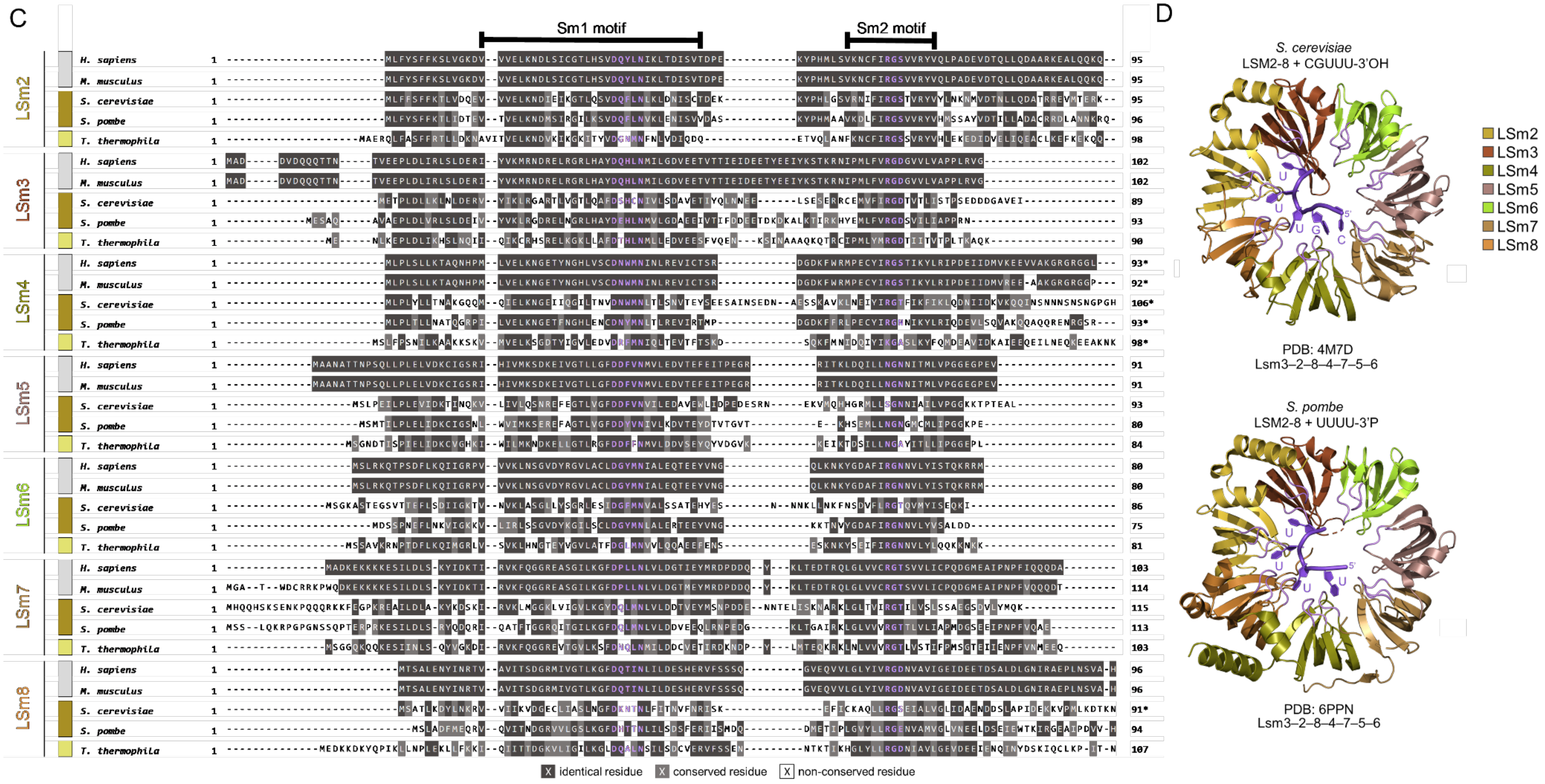


**Figure S8. Primary sequence alignments of the 3'-exonuclease Rex1p, 3'-endonuclease RNase Z / ELAC2 and LSm2-8 complex from different eukaryotic species.**

(A-C) Primary amino acid alignments from different eukaryotic lineages revealing conservation of the 3'-exonuclease Rex1p (A), 3'-endonuclease RNase Z/ELAC2 (B) and LSm2-8 complex (C). Regions important for Rex1p function are highlighted at motif I, motif II and motif III. The RNase Z catalytic domain is shown as PDE. LSm2-8 complex conserved RNA interacting residues are highlighted in purple and conserved Sm1 and Sm2 motifs are shown. A dark grey background indicates identical residues, light grey conserved residues and white indicates no conservation.

(D) High-resolution structures of the Lsm2-8 complex in *Saccharomyces cerevisiae* (PDB: 4M7D) and *Schizosaccharomyces pombe* (PDB: 6PPN) in complex with 3'-uridylate RNA shown in dark purple. The conserved RNA interacting residues, highlighted in purple C, are also shown in light purple and are found in the loops.

**Table S1.** Kd values from EMSAs determining binding of Mlp1 to different processed pre-tRNA intermediates and mature tRNA.

| | **Leu AAG**<br>**$K_d$ (nM)** | **Gln UUA**<br>**$K_d$ (nM)** |
|---|---|---|
| pre-tRNA 5' 3' | 586 ± 15 | 687 ± 44 |
| pre-tRNA 3' | 658 ± 36 | 740 ± 40 |
| pre-tRNA 5' | 714 ± 21 | 804 ± 16 |
| mature tRNA | 831 ± 33 | 801 ± 62 |

**Table S2.** Raw counts for 3'-U ending and 3'-CCA ending tRNAs from raw .fastq files from wild type (WT) input tRNA and Mlp1-immunoprecipitated tRNA (IP). Data was obtained using a custom python script specifically "fishing" sequence specific reads (see Additional Table 2 for fishing sequences) from the raw .fastq file based on the 3'-ends of the tRNA. Data normalization was done by dividing the raw counts by the scaling factor (sum of transcript counts for each replicate divided by 1,000,000), giving counts per million (CPM) values. Fold enrichment of Mlp1-bound mature tRNAs (3'-CCA) and pre-tRNAs (summation of 3'-U) was calculated by taking the ratio of Mlp1-bound tRNA counts and input WT tRNA counts.

| | | pre-tRNA (U-ending) and mature tRNA (CCA-ending) raw counts for WT total tRNAs | | | | | | | | | | | | | | | | | | | | | | | | | | | | | | | | |
|---|---|---|---|---|---|---|---|---|---|---|---|---|---|---|---|---|---|---|---|---|---|---|---|---|---|---|---|---|---|---|---|---|---|---|
| tRNA isotype | tRNA anticodon | CCA_1 | CCA_2 | CCA_3 | U1_1 | U1_2 | U1_3 | U2_1 | U2_2 | U2_3 | U3_1 | U3_2 | U3_3 | U4_1 | U4_2 | U4_3 | U5_1 | U5_2 | U5_3 | U6_1 | U6_2 | U6_3 | U7_1 | U7_2 | U7_3 | U8_1 | U8_2 | U8_3 | U9_1 | U9_2 | U9_3 | U10_1 | U10_2 | U10_3 |
| Ala | AGC | 101773 | 49045 | 61265 | 41 | 30 | 52 | 18 | 5 | 10 | 0 | 1 | 0 | 0 | 0 | 0 | 0 | 0 | 0 | 0 | 0 | 0 | 0 | 0 | 0 | 0 | 0 | 0 | 0 | 0 | 0 | 0 | 0 | 0 |
| Ala | CGC | 17674 | 9944 | 13321 | 9 | 1 | 9 | 1 | 1 | 3 | 0 | 0 | 0 | 0 | 0 | 0 | 0 | 0 | 0 | 0 | 0 | 0 | 0 | 0 | 0 | 0 | 0 | 0 | 0 | 0 | 0 | 0 | 0 | 0 |
| Ala | TGC | 58804 | 25680 | 28108 | 32 | 20 | 48 | 3 | 1 | 4 | 0 | 0 | 1 | 0 | 0 | 0 | 0 | 0 | 0 | 0 | 0 | 0 | 0 | 0 | 0 | 0 | 0 | 0 | 0 | 0 | 0 | 0 | 0 | 0 |
| Arg | ACG | 69845 | 56357 | 77456 | 60 | 56 | 120 | 19 | 6 | 10 | 2 | 7 | 4 | 0 | 0 | 1 | 0 | 0 | 0 | 0 | 0 | 0 | 0 | 0 | 0 | 0 | 0 | 0 | 0 | 0 | 0 | 0 | 0 | 0 |
| Arg | CCG | 1906 | 1188 | 1692 | 1 | 2 | 2 | 0 | 0 | 0 | 0 | 0 | 0 | 0 | 0 | 0 | 0 | 0 | 0 | 0 | 0 | 0 | 0 | 0 | 0 | 0 | 0 | 0 | 0 | 0 | 0 | 0 | 0 | 0 |
| Arg | CCT | 6877 | 5023 | 6250 | 3 | 4 | 10 | 0 | 2 | 0 | 0 | 0 | 0 | 0 | 0 | 0 | 0 | 0 | 0 | 0 | 0 | 0 | 0 | 0 | 0 | 0 | 0 | 0 | 0 | 0 | 0 | 0 | 0 | 0 |
| Arg | TCG | 4727 | 3567 | 5295 | 5 | 8 | 10 | 1 | 0 | 0 | 0 | 0 | 0 | 0 | 0 | 0 | 0 | 0 | 0 | 0 | 0 | 0 | 0 | 0 | 0 | 0 | 0 | 0 | 0 | 0 | 0 | 0 | 0 | 0 |
| Arg | TCT | 655451 | 515155 | 774760 | 718 | 487 | 948 | 51 | 47 | 106 | 16 | 20 | 30 | 7 | 9 | 9 | 0 | 0 | 0 | 0 | 0 | 0 | 0 | 0 | 0 | 0 | 0 | 0 | 0 | 0 | 0 | 0 | 0 | 0 |
| Asn | GTT | 93198 | 47317 | 61766 | 63 | 27 | 41 | 3 | 4 | 1 | 0 | 0 | 0 | 0 | 0 | 0 | 0 | 0 | 0 | 0 | 0 | 0 | 0 | 0 | 0 | 0 | 0 | 0 | 0 | 0 | 0 | 0 | 0 | 0 |
| Asp | GTC | 108197 | 64370 | 83561 | 87 | 67 | 81 | 8 | 23 | 7 | 2 | 1 | 5 | 0 | 0 | 0 | 0 | 0 | 0 | 0 | 0 | 0 | 0 | 0 | 0 | 0 | 0 | 0 | 0 | 0 | 0 | 0 | 0 | 0 |
| Cys | GCA | 123990 | 72413 | 107691 | 119 | 76 | 179 | 4 | 8 | 18 | 0 | 0 | 0 | 0 | 8 | 0 | 4 | 4 | 0 | 0 | 0 | 0 | 0 | 0 | 0 | 0 | 0 | 0 | 0 | 0 | 0 | 0 | 0 | 0 |
| Gln | CTA | 274808 | 148637 | 185931 | 183 | 109 | 190 | 29 | 23 | 22 | 1 | 0 | 10 | 0 | 0 | 6 | 0 | 0 | 0 | 0 | 0 | 0 | 0 | 0 | 0 | 0 | 0 | 0 | 0 | 0 | 0 | 0 | 0 | 0 |
| Gln | TTA | 83090 | 51452 | 64155 | 312 | 180 | 364 | 9 | 1 | 18 | 2 | 0 | 4 | 0 | 0 | 2 | 0 | 0 | 0 | 0 | 0 | 0 | 0 | 0 | 0 | 0 | 0 | 0 | 0 | 0 | 0 | 0 | 0 | 0 |
| Gln | TTG | 589597 | 422248 | 526962 | 490 | 320 | 569 | 66 | 20 | 95 | 24 | 4 | 15 | 7 | 8 | 7 | 0 | 0 | 0 | 0 | 0 | 0 | 0 | 0 | 0 | 0 | 0 | 0 | 0 | 0 | 0 | 0 | 0 | 0 |
| Glu | CTC | 21233 | 12157 | 13254 | 18 | 18 | 23 | 4 | 0 | 11 | 0 | 0 | 0 | 0 | 0 | 0 | 0 | 0 | 0 | 0 | 0 | 0 | 0 | 0 | 0 | 0 | 0 | 0 | 0 | 0 | 0 | 0 | 0 | 0 |
| Glu | TTC | 91903 | 60273 | 70481 | 110 | 117 | 123 | 11 | 14 | 18 | 0 | 0 | 6 | 0 | 0 | 0 | 0 | 0 | 0 | 0 | 0 | 0 | 0 | 0 | 0 | 0 | 0 | 0 | 0 | 0 | 0 | 0 | 0 | 0 |
| Gly | CCC | 2 | 0 | 23 | 0 | 0 | 0 | 0 | 0 | 0 | 0 | 0 | 0 | 0 | 0 | 0 | 0 | 0 | 0 | 0 | 0 | 0 | 0 | 0 | 0 | 0 | 0 | 0 | 0 | 0 | 0 | 0 | 0 | 0 |
| Gly | GCC | 109151 | 49790 | 64024 | 57 | 11 | 44 | 2 | 0 | 2 | 14 | 0 | 3 | 0 | 0 | 0 | 0 | 0 | 0 | 0 | 0 | 0 | 0 | 0 | 0 | 0 | 0 | 0 | 0 | 0 | 0 | 0 | 0 | 0 |
| Gly | TCC | 38765 | 20931 | 32257 | 68 | 22 | 53 | 2 | 0 | 5 | 2 | 0 | 0 | 0 | 0 | 0 | 0 | 0 | 0 | 0 | 0 | 0 | 0 | 0 | 0 | 0 | 0 | 0 | 0 | 0 | 0 | 0 | 0 | 0 |
| His | GTG | 46189 | 28199 | 28886 | 23 | 18 | 19 | 1 | 1 | 9 | 0 | 0 | 0 | 0 | 8 | 0 | 0 | 0 | 0 | 0 | 0 | 0 | 0 | 0 | 0 | 0 | 0 | 0 | 0 | 0 | 0 | 0 | 0 | 0 |
| Ile | AAT | 194897 | 139549 | 167646 | 123 | 53 | 113 | 23 | 12 | 5 | 0 | 0 | 0 | 0 | 0 | 3 | 0 | 0 | 0 | 0 | 0 | 0 | 0 | 0 | 0 | 0 | 0 | 0 | 0 | 0 | 0 | 0 | 0 | 0 |
| Ile | TAT | 25298 | 16946 | 19701 | 27 | 10 | 46 | 0 | 0 | 6 | 0 | 6 | 0 | 0 | 0 | 0 | 0 | 0 | 0 | 0 | 0 | 0 | 0 | 0 | 0 | 0 | 0 | 0 | 0 | 0 | 0 | 0 | 0 | 0 |
| Leu | AAG | 736823 | 371241 | 387961 | 372 | 230 | 274 | 38 | 36 | 10 | 0 | 0 | 8 | 0 | 0 | 0 | 0 | 0 | 0 | 0 | 0 | 0 | 2 | 0 | 0 | 0 | 0 | 0 | 0 | 0 | 0 | 0 | 0 | 0 |
| Leu | CAA | 534721 | 385328 | 330488 | 673 | 587 | 587 | 259 | 130 | 206 | 38 | 5 | 15 | 4 | 3 | 8 | 2 | 0 | 0 | 0 | 0 | 0 | 0 | 0 | 0 | 0 | 0 | 0 | 0 | 0 | 0 | 0 | 0 | 0 |
| Leu | CAG | 694731 | 350058 | 366549 | 287 | 174 | 207 | 33 | 35 | 8 | 0 | 0 | 8 | 0 | 0 | 0 | 0 | 0 | 0 | 0 | 0 | 0 | 2 | 0 | 0 | 0 | 0 | 0 | 0 | 0 | 0 | 0 | 0 | 0 |
| Leu | TAA | 81109 | 57291 | 76435 | 151 | 84 | 151 | 17 | 5 | 18 | 0 | 0 | 1 | 0 | 0 | 0 | 0 | 0 | 0 | 0 | 0 | 0 | 0 | 0 | 0 | 0 | 0 | 0 | 0 | 0 | 0 | 0 | 0 | 0 |
| Leu | TAG | 54400 | 31709 | 35419 | 24 | 13 | 20 | 5 | 2 | 0 | 7 | 0 | 0 | 0 | 0 | 4 | 3 | 0 | 0 | 0 | 0 | 0 | 0 | 0 | 0 | 0 | 0 | 0 | 0 | 0 | 0 | 0 | 0 | 0 |
| Lys | TTT | 81449 | 40102 | 47166 | 27 | 47 | 32 | 3 | 2 | 4 | 0 | 0 | 0 | 0 | 0 | 0 | 0 | 0 | 0 | 0 | 0 | 0 | 0 | 0 | 0 | 0 | 0 | 0 | 0 | 0 | 0 | 0 | 0 | 0 |
| Met | CAT | 122134 | 83729 | 115064 | 176 | 119 | 213 | 9 | 11 | 19 | 2 | 0 | 1 | 0 | 1 | 1 | 0 | 0 | 0 | 0 | 0 | 0 | 0 | 0 | 0 | 0 | 0 | 0 | 0 | 0 | 0 | 0 | 0 | 0 |
| Phe | AAA | 0 | 0 | 0 | 0 | 0 | 0 | 0 | 0 | 0 | 0 | 0 | 0 | 0 | 0 | 0 | 0 | 0 | 0 | 0 | 0 | 0 | 0 | 0 | 0 | 0 | 0 | 0 | 0 | 0 | 0 | 0 | 0 | 0 |
| Phe | GAA | 73674 | 50329 | 56535 | 52 | 42 | 31 | 4 | 5 | 3 | 0 | 0 | 10 | 0 | 0 | 8 | 0 | 0 | 0 | 0 | 0 | 0 | 0 | 0 | 0 | 0 | 0 | 0 | 0 | 0 | 0 | 0 | 0 | 0 |
| Pro | AGG | 64362 | 35241 | 51175 | 131 | 49 | 118 | 24 | 7 | 17 | 2 | 0 | 1 | 2 | 0 | 0 | 0 | 0 | 0 | 0 | 0 | 0 | 0 | 0 | 0 | 0 | 0 | 0 | 0 | 0 | 0 | 0 | 0 | 0 |
| Pro | CGG | 4039 | 1924 | 2166 | 3 | 0 | 1 | 0 | 0 | 0 | 0 | 0 | 0 | 0 | 0 | 0 | 0 | 0 | 0 | 0 | 0 | 0 | 0 | 0 | 0 | 0 | 0 | 0 | 0 | 0 | 0 | 0 | 0 | 0 |
| Pro | TGG | 19765 | 10741 | 15525 | 42 | 33 | 27 | 0 | 2 | 2 | 0 | 0 | 3 | 0 | 0 | 2 | 0 | 0 | 0 | 0 | 0 | 0 | 0 | 0 | 0 | 0 | 0 | 0 | 0 | 0 | 0 | 0 | 0 | 0 |
| Ser | AGA | 228626 | 145814 | 205261 | 239 | 110 | 306 | 47 | 25 | 81 | 6 | 0 | 4 | 0 | 0 | 3 | 0 | 0 | 0 | 0 | 0 | 0 | 0 | 0 | 0 | 0 | 0 | 0 | 0 | 0 | 0 | 0 | 0 | 0 |
| Ser | CGA | 19689 | 17749 | 21887 | 25 | 14 | 39 | 3 | 0 | 0 | 0 | 0 | 0 | 0 | 0 | 0 | 0 | 0 | 0 | 0 | 0 | 0 | 0 | 0 | 0 | 0 | 0 | 0 | 0 | 0 | 0 | 0 | 0 | 0 |
| Ser | GCT | 83449 | 46174 | 61004 | 60 | 24 | 54 | 6 | 2 | 0 | 9 | 0 | 0 | 0 | 0 | 0 | 0 | 0 | 0 | 3 | 0 | 0 | 0 | 0 | 0 | 0 | 0 | 0 | 0 | 0 | 0 | 0 | 0 | 0 |
| Ser | TGA | 74794 | 52775 | 69731 | 48 | 16 | 36 | 9 | 0 | 13 | 0 | 0 | 1 | 3 | 0 | 0 | 0 | 0 | 0 | 0 | 0 | 0 | 0 | 0 | 0 | 0 | 0 | 0 | 0 | 0 | 0 | 0 | 0 | 0 |
| Thr | AGT | 76120 | 43784 | 53703 | 76 | 29 | 121 | 9 | 14 | 13 | 0 | 0 | 0 | 0 | 0 | 0 | 3 | 0 | 0 | 0 | 0 | 0 | 0 | 0 | 0 | 0 | 0 | 0 | 0 | 0 | 0 | 0 | 0 | 0 |
| Thr | CGT | 3019 | 2628 | 2513 | 1 | 2 | 3 | 0 | 0 | 0 | 0 | 0 | 0 | 0 | 0 | 0 | 0 | 0 | 0 | 0 | 0 | 0 | 0 | 0 | 0 | 0 | 0 | 0 | 0 | 0 | 0 | 0 | 0 | 0 |
| Thr | TGT | 226719 | 90263 | 66532 | 99 | 45 | 38 | 4 | 2 | 1 | 0 | 0 | 0 | 0 | 0 | 0 | 0 | 0 | 0 | 0 | 0 | 0 | 0 | 0 | 0 | 0 | 0 | 0 | 0 | 0 | 0 | 0 | 0 | 0 |
| Trp | CCA | 50436 | 29034 | 36424 | 49 | 62 | 35 | 2 | 5 | 1 | 0 | 0 | 0 | 0 | 0 | 0 | 0 | 0 | 0 | 0 | 0 | 0 | 0 | 0 | 0 | 0 | 0 | 0 | 0 | 0 | 0 | 0 | 0 | 0 |
| Tyr | GTA | 11817 | 9834 | 11524 | 27 | 37 | 82 | 20 | 26 | 79 | 31 | 7 | 11 | 0 | 0 | 0 | 0 | 0 | 0 | 0 | 0 | 0 | 0 | 0 | 0 | 0 | 0 | 0 | 0 | 0 | 0 | 0 | 0 | 0 |
| Val | AAC | 43617 | 21142 | 23822 | 32 | 21 | 47 | 4 | 0 | 4 | 0 | 0 | 0 | 0 | 0 | 0 | 0 | 0 | 0 | 0 | 0 | 0 | 0 | 0 | 0 | 0 | 0 | 0 | 0 | 0 | 0 | 0 | 0 | 0 |
| Val | CAC | 6095 | 2944 | 4309 | 35 | 12 | 69 | 0 | 0 | 0 | 0 | 0 | 0 | 0 | 0 | 0 | 0 | 0 | 0 | 0 | 0 | 0 | 0 | 0 | 0 | 0 | 0 | 0 | 0 | 0 | 0 | 0 | 0 | 0 |
| Val | TAC | 47244 | 23873 | 28946 | 35 | 15 | 29 | 4 | 1 | 1 | 0 | 0 | 0 | 0 | 0 | 0 | 0 | 0 | 0 | 0 | 0 | 0 | 0 | 0 | 0 | 0 | 0 | 0 | 0 | 0 | 0 | 0 | 0 | 0 |

| | | premature (U-ending) and mature (CCA-ending) tRNA counts for MLP1-immunoprecipitated tRNAs | | | | | | | | | | | | | | | | | | | | | | | | | | | | | | | | |
|---|---|---|---|---|---|---|---|---|---|---|---|---|---|---|---|---|---|---|---|---|---|---|---|---|---|---|---|---|---|---|---|---|---|---|
| **tRNA isotype** | **tRNA anticodon** | **CCA_1** | **CCA_2** | **CCA_3** | **U1_1** | **U1_2** | **U1_3** | **U2_1** | **U2_2** | **U2_3** | **U3_1** | **U3_2** | **U3_3** | **U4_1** | **U4_2** | **U4_3** | **U5_1** | **U5_2** | **U5_3** | **U6_1** | **U6_2** | **U6_3** | **U7_1** | **U7_2** | **U7_3** | **U8_1** | **U8_2** | **U8_3** | **U9_1** | **U9_2** | **U9_3** | **U10_1** | **U10_2** | **U10_3** |
| Ala | AGC | 12790 | 18607 | 13616 | 27 | 19 | 7 | 3 | 0 | 0 | 13 | 0 | 0 | 0 | 0 | 0 | 0 | 0 | 0 | 0 | 0 | 0 | 0 | 0 | 0 | 0 | 0 | 0 | 0 | 0 | 0 | 0 | 0 | 0 |
| Ala | CGC | 1807 | 2306 | 2050 | 0 | 0 | 9 | 0 | 0 | 0 | 0 | 0 | 0 | 0 | 0 | 0 | 0 | 0 | 0 | 0 | 0 | 0 | 0 | 0 | 0 | 0 | 0 | 0 | 0 | 0 | 0 | 0 | 0 | 0 |
| Ala | TGC | 7717 | 10867 | 10919 | 4 | 2 | 30 | 0 | 0 | 0 | 0 | 0 | 0 | 0 | 0 | 0 | 0 | 0 | 0 | 0 | 0 | 0 | 0 | 0 | 0 | 0 | 0 | 0 | 0 | 0 | 0 | 0 | 0 | 0 |
| Arg | ACG | 12057 | 14757 | 16553 | 27 | 27 | 39 | 0 | 1 | 4 | 0 | 0 | 0 | 0 | 0 | 0 | 0 | 0 | 0 | 0 | 0 | 0 | 0 | 0 | 0 | 0 | 0 | 0 | 0 | 0 | 0 | 0 | 0 | 0 |
| Arg | CCG | 485 | 734 | 684 | 1 | 1 | 0 | 0 | 0 | 0 | 0 | 0 | 0 | 0 | 0 | 0 | 0 | 0 | 0 | 0 | 0 | 0 | 0 | 0 | 0 | 0 | 0 | 0 | 0 | 0 | 0 | 0 | 0 | 0 |
| Arg | CCT | 3471 | 4118 | 2867 | 4 | 0 | 2 | 0 | 2 | 12 | 0 | 0 | 0 | 0 | 0 | 0 | 0 | 0 | 0 | 0 | 0 | 0 | 0 | 0 | 0 | 0 | 0 | 0 | 0 | 0 | 0 | 0 | 0 | 0 |
| Arg | TCG | 1282 | 1895 | 1583 | 2 | 4 | 11 | 0 | 0 | 0 | 0 | 0 | 0 | 0 | 0 | 0 | 0 | 0 | 0 | 0 | 0 | 0 | 0 | 0 | 0 | 0 | 0 | 0 | 0 | 0 | 0 | 0 | 0 | 0 |
| Arg | TCT | 121075 | 172409 | 153047 | 82 | 125 | 117 | 4 | 31 | 2 | 27 | 29 | 32 | 0 | 0 | 0 | 0 | 0 | 0 | 0 | 0 | 0 | 0 | 0 | 0 | 0 | 0 | 0 | 0 | 0 | 0 | 0 | 0 | 0 |
| Asn | GTT | 24429 | 36169 | 28634 | 12 | 34 | 27 | 0 | 17 | 2 | 12 | 22 | 0 | 0 | 0 | 0 | 0 | 0 | 0 | 0 | 0 | 0 | 0 | 0 | 0 | 0 | 0 | 0 | 0 | 0 | 0 | 0 | 0 | 0 |
| Asp | GTC | 33001 | 45745 | 35673 | 21 | 52 | 43 | 64 | 45 | 71 | 16 | 16 | 23 | 0 | 0 | 7 | 0 | 0 | 1 | 0 | 0 | 0 | 0 | 0 | 0 | 0 | 0 | 0 | 0 | 0 | 0 | 0 | 0 | 0 |
| Cys | GCA | 12559 | 20982 | 18480 | 30 | 15 | 8 | 16 | 18 | 31 | 26 | 16 | 52 | 0 | 3 | 17 | 0 | 0 | 0 | 0 | 0 | 0 | 0 | 0 | 0 | 0 | 0 | 0 | 0 | 0 | 0 | 0 | 0 | 0 |
| Gln | CTA | 20308 | 31636 | 27251 | 12 | 33 | 43 | 15 | 3 | 10 | 33 | 76 | 101 | 7 | 0 | 23 | 0 | 0 | 0 | 0 | 0 | 0 | 0 | 0 | 0 | 0 | 0 | 0 | 0 | 0 | 0 | 0 | 0 | 0 |
| Gln | TTA | 4181 | 6675 | 5297 | 15 | 100 | 115 | 6 | 17 | 44 | 17 | 17 | 29 | 0 | 0 | 0 | 0 | 0 | 0 | 0 | 0 | 0 | 0 | 0 | 0 | 0 | 0 | 0 | 0 | 0 | 0 | 0 | 0 | 0 |
| Gln | TTG | 83793 | 115812 | 107411 | 40 | 104 | 60 | 6 | 6 | 3 | 15 | 0 | 37 | 0 | 0 | 0 | 0 | 0 | 0 | 0 | 0 | 0 | 0 | 0 | 0 | 0 | 0 | 0 | 0 | 0 | 0 | 0 | 0 | 0 |
| Glu | CTC | 2810 | 4166 | 4151 | 9 | 3 | 6 | 0 | 0 | 22 | 0 | 0 | 0 | 0 | 0 | 0 | 0 | 0 | 0 | 0 | 0 | 0 | 0 | 0 | 0 | 0 | 0 | 0 | 0 | 0 | 0 | 0 | 0 | 0 |
| Glu | TTC | 26920 | 33372 | 32085 | 99 | 122 | 104 | 25 | 68 | 81 | 30 | 30 | 10 | 0 | 1 | 0 | 0 | 0 | 1 | 0 | 0 | 0 | 0 | 0 | 0 | 0 | 0 | 0 | 0 | 0 | 0 | 0 | 0 | 0 |
| Gly | CCC | 0 | 0 | 14 | 0 | 0 | 0 | 0 | 0 | 0 | 0 | 0 | 0 | 0 | 0 | 0 | 0 | 0 | 0 | 0 | 0 | 0 | 0 | 0 | 0 | 0 | 0 | 0 | 0 | 0 | 0 | 0 | 0 | 0 |
| Gly | GCC | 47487 | 63069 | 52703 | 9 | 18 | 30 | 0 | 0 | 0 | 0 | 56 | 0 | 21 | 63 | 19 | 0 | 0 | 0 | 0 | 0 | 0 | 0 | 0 | 0 | 0 | 0 | 0 | 0 | 0 | 0 | 0 | 0 | 0 |
| Gly | TCC | 4397 | 6995 | 5856 | 42 | 13 | 42 | 25 | 21 | 24 | 16 | 18 | 0 | 5 | 0 | 4 | 0 | 0 | 0 | 0 | 0 | 0 | 0 | 0 | 0 | 0 | 0 | 0 | 0 | 0 | 0 | 0 | 0 | 0 |
| His | GTG | 18117 | 25645 | 17210 | 2 | 6 | 19 | 28 | 26 | 29 | 0 | 19 | 45 | 0 | 0 | 0 | 0 | 0 | 0 | 0 | 0 | 0 | 0 | 0 | 0 | 0 | 0 | 0 | 0 | 0 | 0 | 0 | 0 | 0 |
| Ile | AAT | 58883 | 105760 | 71718 | 24 | 71 | 36 | 2 | 2 | 0 | 0 | 24 | 1 | 13 | 2 | 9 | 0 | 0 | 0 | 0 | 0 | 0 | 0 | 0 | 0 | 0 | 0 | 0 | 0 | 0 | 0 | 0 | 0 | 0 |
| Ile | TAT | 3852 | 6245 | 5081 | 26 | 2 | 34 | 0 | 31 | 7 | 0 | 1 | 0 | 27 | 32 | 0 | 14 | 0 | 4 | 11 | 0 | 0 | 0 | 0 | 0 | 0 | 0 | 0 | 0 | 0 | 0 | 0 | 0 | 0 |
| Leu | AAG | 303102 | 366843 | 206948 | 148 | 186 | 127 | 38 | 74 | 35 | 23 | 1 | 55 | 80 | 39 | 23 | 0 | 22 | 0 | 17 | 1 | 0 | 0 | 0 | 1 | 0 | 5 | 31 | 0 | 0 | 0 | 0 | 0 | 0 |
| Leu | CAA | 162010 | 209902 | 143378 | 56 | 88 | 94 | 74 | 15 | 38 | 0 | 3 | 5 | 0 | 0 | 12 | 0 | 1 | 1 | 0 | 0 | 0 | 0 | 0 | 0 | 0 | 0 | 0 | 0 | 0 | 0 | 0 | 0 | 0 |
| Leu | CAG | 284604 | 345178 | 194258 | 112 | 142 | 90 | 38 | 71 | 3 | 22 | 1 | 52 | 80 | 39 | 21 | 0 | 20 | 0 | 14 | 1 | 0 | 0 | 0 | 1 | 0 | 4 | 31 | 0 | 0 | 0 | 0 | 0 | 0 |
| Leu | TAA | 13122 | 16157 | 15068 | 65 | 47 | 26 | 24 | 25 | 5 | 0 | 0 | 8 | 0 | 14 | 31 | 0 | 26 | 0 | 0 | 0 | 0 | 0 | 0 | 0 | 0 | 0 | 0 | 0 | 0 | 0 | 0 | 0 | 0 |
| Leu | TAG | 26395 | 30450 | 22376 | 7 | 37 | 47 | 2 | 1 | 1 | 26 | 24 | 27 | 0 | 84 | 30 | 0 | 0 | 7 | 0 | 0 | 0 | 0 | 0 | 0 | 0 | 0 | 0 | 0 | 0 | 0 | 0 | 0 | 0 |
| Lys | TTT | 19857 | 13620 | 10960 | 35 | 54 | 32 | 14 | 0 | 0 | 28 | 0 | 18 | 72 | 0 | 30 | 12 | 2 | 1 | 0 | 31 | 0 | 0 | 0 | 0 | 18 | 0 | 7 | 0 | 0 | 0 | 0 | 0 | 0 |
| Met | CAT | 27999 | 43139 | 33348 | 131 | 236 | 262 | 23 | 38 | 15 | 8 | 13 | 5 | 17 | 0 | 0 | 0 | 0 | 0 | 0 | 0 | 0 | 0 | 0 | 0 | 0 | 0 | 0 | 0 | 0 | 0 | 0 | 0 | 0 |
| Phe | AAA | 12421 | 18200 | 14340 | 172 | 376 | 262 | 50 | 2 | 34 | 0 | 0 | 28 | 0 | 0 | 0 | 0 | 0 | 0 | 0 | 2 | 0 | 0 | 0 | 0 | 0 | 0 | 0 | 0 | 0 | 0 | 0 | 0 | 0 |
| Phe | GAA | 16540 | 23446 | 15822 | 68 | 65 | 33 | 23 | 45 | 9 | 1 | 0 | 7 | 16 | 0 | 17 | 0 | 0 | 0 | 0 | 0 | 0 | 0 | 18 | 0 | 0 | 0 | 0 | 0 | 10 | 0 | 0 | 0 | 0 |
| Pro | AGG | 0 | 0 | 0 | 0 | 0 | 0 | 0 | 0 | 0 | 0 | 0 | 0 | 0 | 0 | 0 | 0 | 0 | 0 | 0 | 0 | 0 | 0 | 0 | 0 | 0 | 0 | 0 | 0 | 0 | 0 | 0 | 0 | 0 |
| Pro | CGG | 1214 | 1558 | 1427 | 0 | 0 | 1 | 0 | 0 | 0 | 0 | 0 | 0 | 0 | 0 | 0 | 0 | 0 | 0 | 0 | 0 | 0 | 0 | 0 | 0 | 0 | 0 | 0 | 0 | 0 | 0 | 0 | 0 | 0 |
| Pro | TGG | 5158 | 8927 | 6768 | 23 | 31 | 15 | 0 | 0 | 1 | 9 | 16 | 0 | 13 | 8 | 13 | 0 | 0 | 0 | 0 | 0 | 0 | 0 | 0 | 0 | 0 | 0 | 0 | 0 | 0 | 0 | 0 | 0 | 0 |
| Ser | AGA | 45732 | 57271 | 42418 | 130 | 47 | 79 | 37 | 16 | 1 | 140 | 135 | 19 | 668 | 225 | 129 | 156 | 16 | 25 | 7 | 10 | 0 | 11 | 0 | 0 | 0 | 0 | 0 | 0 | 0 | 0 | 0 | 11 | 0 |
| Ser | CGA | 6195 | 7190 | 8046 | 224 | 111 | 219 | 711 | 472 | 274 | 0 | 0 | 20 | 0 | 26 | 6 | 9 | 7 | 0 | 11 | 0 | 0 | 0 | 0 | 0 | 0 | 0 | 0 | 0 | 0 | 0 | 0 | 0 | 0 |
| Ser | GCT | 18535 | 28469 | 19044 | 148 | 70 | 146 | 137 | 119 | 188 | 44 | 13 | 7 | 12 | 0 | 54 | 0 | 69 | 3 | 0 | 63 | 0 | 0 | 15 | 11 | 0 | 0 | 0 | 0 | 0 | 0 | 0 | 0 | 0 |
| Ser | TGA | 12304 | 17710 | 14669 | 47 | 5 | 9 | 74 | 39 | 9 | 222 | 226 | 162 | 1 | 1 | 1 | 0 | 0 | 0 | 0 | 0 | 0 | 0 | 0 | 0 | 0 | 0 | 6 | 0 | 0 | 0 | 0 | 0 | 0 |
| Thr | AGT | 12275 | 18076 | 12647 | 2 | 22 | 26 | 15 | 0 | 9 | 0 | 0 | 16 | 0 | 0 | 0 | 0 | 0 | 0 | 0 | 0 | 0 | 0 | 0 | 0 | 0 | 0 | 0 | 0 | 0 | 0 | 0 | 0 | 0 |
| Thr | CGT | 1484 | 2567 | 1843 | 3 | 23 | 0 | 38 | 1 | 0 | 0 | 0 | 0 | 0 | 0 | 0 | 0 | 0 | 1 | 0 | 0 | 0 | 0 | 0 | 0 | 6 | 17 | 0 | 0 | 0 | 0 | 0 | 0 | 0 |
| Thr | TGT | 55865 | 91243 | 60231 | 16 | 36 | 28 | 0 | 0 | 0 | 0 | 1 | 0 | 0 | 0 | 0 | 0 | 0 | 0 | 0 | 0 | 0 | 0 | 0 | 0 | 0 | 0 | 0 | 0 | 0 | 0 | 0 | 0 | 0 |
| Trp | CCA | 8134 | 11600 | 11939 | 184 | 46 | 129 | 12 | 0 | 19 | 0 | 0 | 25 | 0 | 12 | 6 | 0 | 0 | 0 | 0 | 0 | 0 | 0 | 0 | 0 | 0 | 0 | 0 | 0 | 0 | 0 | 0 | 0 | 0 |
| Tyr | GTA | 3551 | 5553 | 4597 | 12 | 5 | 23 | 100 | 205 | 251 | 136 | 175 | 239 | 9 | 1 | 7 | 0 | 0 | 0 | 25 | 0 | 0 | 0 | 0 | 13 | 9 | 33 | 12 | 0 | 0 | 0 | 0 | 0 | 0 |
| Val | AAC | 7018 | 8275 | 6321 | 93 | 125 | 55 | 1 | 0 | 0 | 0 | 0 | 0 | 0 | 0 | 0 | 1 | 0 | 0 | 0 | 0 | 0 | 0 | 0 | 0 | 0 | 0 | 0 | 0 | 0 | 0 | 0 | 0 | 0 |
| Val | CAC | 1399 | 1881 | 1464 | 5 | 0 | 52 | 0 | 0 | 1 | 9 | 34 | 18 | 0 | 1 | 9 | 0 | 0 | 0 | 0 | 0 | 0 | 0 | 0 | 0 | 0 | 0 | 0 | 0 | 0 | 0 | 0 | 0 | 0 |
| Val | TAC | 5806 | 7978 | 5496 | 1 | 54 | 21 | 0 | 0 | 0 | 0 | 0 | 0 | 0 | 0 | 0 | 0 | 0 | 0 | 0 | 0 | 0 | 0 | 0 | 0 | 0 | 0 | 0 | 0 | 0 | 0 | 0 | 0 | 0 |

**Table S3.** Raw counts for 3'-U ending and 3'-CCA ending tRNAs from raw .fastq files. Data was obtained using a custom python script specifically "fishing" sequence specific reads (see Additional Table 2 for fishing sequences) from the raw .fastq file based on the 3'-ends of the tRNA. Data normalization was done by dividing the raw counts by the scaling factor (sum of transcript counts for each replicate divided by 1,000,000), giving counts per million (CPM) values. The average log10 value of the cumulative TPM is shown in the heatmap in Figure 5C.

| | | premature (U-ending) and mature (CCA-ending) tRNA counts for wild type (WT) total tRNAs | | | | | | | | | | | | | | | | | | | | | | | | | | | | | | | | |
|---|---|---|---|---|---|---|---|---|---|---|---|---|---|---|---|---|---|---|---|---|---|---|---|---|---|---|---|---|---|---|---|---|---|---|
| tRNA isotype | tRNA anticodon | CCA_1 | CCA_2 | CCA_3 | U1_1 | U1_2 | U1_3 | U2_1 | U2_2 | U2_3 | U3_1 | U3_2 | U3_3 | U4_1 | U4_2 | U4_3 | U5_1 | U5_2 | U5_3 | U6_1 | U6_2 | U6_3 | U7_1 | U7_2 | U7_3 | U8_1 | U8_2 | U8_3 | U9_1 | U9_2 | U9_3 | U10_1 | U10_2 | U10_3 |
| Ala | AGC | 101773 | 49045 | 61265 | 41 | 30 | 52 | 18 | 5 | 10 | 0 | 1 | 0 | 0 | 0 | 0 | 0 | 0 | 0 | 0 | 0 | 0 | 0 | 0 | 0 | 0 | 0 | 0 | 0 | 0 | 0 | 0 | 0 | 0 |
| Ala | CGC | 17674 | 9944 | 13321 | 9 | 1 | 9 | 1 | 1 | 3 | 0 | 0 | 0 | 0 | 0 | 0 | 0 | 0 | 0 | 0 | 0 | 0 | 0 | 0 | 0 | 0 | 0 | 0 | 0 | 0 | 0 | 0 | 0 | 0 |
| Ala | TGC | 58804 | 25680 | 28108 | 32 | 20 | 48 | 3 | 1 | 4 | 0 | 0 | 1 | 0 | 0 | 0 | 0 | 0 | 0 | 0 | 0 | 0 | 0 | 0 | 0 | 0 | 0 | 0 | 0 | 0 | 0 | 0 | 0 | 0 |
| Arg | ACG | 69845 | 56357 | 77456 | 60 | 56 | 120 | 19 | 6 | 10 | 2 | 7 | 4 | 0 | 0 | 1 | 0 | 0 | 0 | 0 | 0 | 0 | 0 | 0 | 0 | 0 | 0 | 0 | 0 | 0 | 0 | 0 | 0 | 0 |
| Arg | CCG | 1906 | 1188 | 1692 | 1 | 2 | 2 | 0 | 0 | 0 | 0 | 0 | 0 | 0 | 0 | 0 | 0 | 0 | 0 | 0 | 0 | 0 | 0 | 0 | 0 | 0 | 0 | 0 | 0 | 0 | 0 | 0 | 0 | 0 |
| Arg | CCT | 6877 | 5023 | 6250 | 3 | 4 | 10 | 0 | 2 | 0 | 0 | 0 | 0 | 0 | 0 | 0 | 0 | 0 | 0 | 0 | 0 | 0 | 0 | 0 | 0 | 0 | 0 | 0 | 0 | 0 | 0 | 0 | 0 | 0 |
| Arg | TCG | 4727 | 3567 | 5295 | 5 | 8 | 10 | 1 | 0 | 0 | 0 | 0 | 0 | 0 | 0 | 0 | 0 | 0 | 0 | 0 | 0 | 0 | 0 | 0 | 0 | 0 | 0 | 0 | 0 | 0 | 0 | 0 | 0 | 0 |
| Arg | TCT | 655451 | 515155 | 774760 | 718 | 487 | 948 | 51 | 47 | 106 | 16 | 20 | 30 | 7 | 9 | 9 | 0 | 0 | 0 | 0 | 0 | 0 | 0 | 0 | 0 | 0 | 0 | 0 | 0 | 0 | 0 | 0 | 0 | 0 |
| Asn | GTT | 93198 | 47317 | 61766 | 63 | 27 | 41 | 3 | 4 | 1 | 0 | 0 | 0 | 0 | 0 | 0 | 0 | 0 | 0 | 0 | 0 | 0 | 0 | 0 | 0 | 0 | 0 | 0 | 0 | 0 | 0 | 0 | 0 | 0 |
| Asp | GTC | 108197 | 64370 | 83561 | 87 | 67 | 81 | 8 | 23 | 7 | 2 | 1 | 5 | 0 | 0 | 0 | 0 | 0 | 0 | 0 | 0 | 0 | 0 | 0 | 0 | 0 | 0 | 0 | 0 | 0 | 0 | 0 | 0 | 0 |
| Cys | GCA | 123990 | 72413 | 107691 | 119 | 76 | 179 | 4 | 8 | 18 | 0 | 0 | 0 | 0 | 8 | 0 | 4 | 4 | 0 | 0 | 0 | 0 | 0 | 0 | 0 | 0 | 0 | 0 | 0 | 0 | 0 | 0 | 0 | 0 |
| Gln | CTA | 274808 | 148637 | 185931 | 183 | 109 | 190 | 29 | 23 | 22 | 1 | 0 | 10 | 0 | 0 | 6 | 0 | 0 | 0 | 0 | 0 | 0 | 0 | 0 | 0 | 0 | 0 | 0 | 0 | 0 | 0 | 0 | 0 | 0 |
| Gln | TTA | 83090 | 51452 | 64155 | 312 | 180 | 364 | 9 | 1 | 18 | 2 | 0 | 4 | 0 | 0 | 2 | 0 | 0 | 0 | 0 | 0 | 0 | 0 | 0 | 0 | 0 | 0 | 0 | 0 | 0 | 0 | 0 | 0 | 0 |
| Gln | TTG | 589597 | 422248 | 526962 | 490 | 320 | 569 | 66 | 20 | 95 | 24 | 4 | 15 | 7 | 8 | 7 | 0 | 0 | 0 | 0 | 0 | 0 | 0 | 0 | 0 | 0 | 0 | 0 | 0 | 0 | 0 | 0 | 0 | 0 |
| Glu | CTC | 21233 | 12157 | 13254 | 18 | 18 | 23 | 4 | 0 | 11 | 0 | 0 | 0 | 0 | 0 | 0 | 0 | 0 | 0 | 0 | 0 | 0 | 0 | 0 | 0 | 0 | 0 | 0 | 0 | 0 | 0 | 0 | 0 | 0 |
| Glu | TTC | 91903 | 60273 | 70481 | 110 | 117 | 123 | 11 | 14 | 18 | 0 | 0 | 6 | 0 | 0 | 0 | 0 | 0 | 0 | 0 | 0 | 0 | 0 | 0 | 0 | 0 | 0 | 0 | 0 | 0 | 0 | 0 | 0 | 0 |
| Gly | CCC | 2 | 0 | 23 | 0 | 0 | 0 | 0 | 0 | 0 | 0 | 0 | 0 | 0 | 0 | 0 | 0 | 0 | 0 | 0 | 0 | 0 | 0 | 0 | 0 | 0 | 0 | 0 | 0 | 0 | 0 | 0 | 0 | 0 |
| Gly | GCC | 109151 | 49790 | 64024 | 57 | 11 | 44 | 2 | 0 | 2 | 14 | 0 | 3 | 0 | 0 | 0 | 0 | 0 | 0 | 0 | 0 | 0 | 0 | 0 | 0 | 0 | 0 | 0 | 0 | 0 | 0 | 0 | 0 | 0 |
| Gly | TCC | 38765 | 20931 | 32257 | 68 | 22 | 53 | 2 | 0 | 5 | 2 | 0 | 0 | 0 | 0 | 0 | 0 | 0 | 0 | 0 | 0 | 0 | 0 | 0 | 0 | 0 | 0 | 0 | 0 | 0 | 0 | 0 | 0 | 0 |
| His | GTG | 46189 | 28199 | 28886 | 23 | 18 | 19 | 1 | 1 | 9 | 0 | 0 | 0 | 0 | 8 | 0 | 0 | 0 | 0 | 0 | 0 | 0 | 0 | 0 | 0 | 0 | 0 | 0 | 0 | 0 | 0 | 0 | 0 | 0 |
| Ile | AAT | 194897 | 139549 | 167646 | 123 | 53 | 113 | 23 | 12 | 5 | 0 | 0 | 0 | 0 | 0 | 3 | 0 | 0 | 0 | 0 | 0 | 0 | 0 | 0 | 0 | 0 | 0 | 0 | 0 | 0 | 0 | 0 | 0 | 0 |
| Ile | TAT | 25298 | 16946 | 19701 | 27 | 10 | 46 | 0 | 0 | 6 | 0 | 6 | 0 | 0 | 0 | 0 | 0 | 0 | 0 | 0 | 0 | 0 | 0 | 0 | 0 | 0 | 0 | 0 | 0 | 0 | 0 | 0 | 0 | 0 |
| Leu | AAG | 736823 | 371241 | 387961 | 372 | 230 | 274 | 38 | 36 | 10 | 0 | 0 | 8 | 0 | 0 | 0 | 0 | 0 | 0 | 0 | 0 | 0 | 2 | 0 | 0 | 0 | 0 | 0 | 0 | 0 | 0 | 0 | 0 | 0 |
| Leu | CAA | 534721 | 385328 | 330488 | 673 | 587 | 587 | 259 | 130 | 206 | 38 | 5 | 15 | 4 | 3 | 8 | 2 | 0 | 0 | 0 | 0 | 0 | 0 | 0 | 0 | 0 | 0 | 0 | 0 | 0 | 0 | 0 | 0 | 0 |
| Leu | CAG | 694731 | 350058 | 366549 | 287 | 174 | 207 | 33 | 35 | 8 | 0 | 0 | 8 | 0 | 0 | 0 | 0 | 0 | 0 | 0 | 0 | 0 | 2 | 0 | 0 | 0 | 0 | 0 | 0 | 0 | 0 | 0 | 0 | 0 |
| Leu | TAA | 81109 | 57291 | 76435 | 151 | 84 | 151 | 17 | 5 | 18 | 0 | 0 | 1 | 0 | 0 | 0 | 0 | 0 | 0 | 0 | 0 | 0 | 0 | 0 | 0 | 0 | 0 | 0 | 0 | 0 | 0 | 0 | 0 | 0 |
| Leu | TAG | 54400 | 31709 | 35419 | 24 | 13 | 20 | 5 | 2 | 0 | 7 | 0 | 0 | 0 | 0 | 4 | 3 | 0 | 0 | 0 | 0 | 0 | 0 | 0 | 0 | 0 | 0 | 0 | 0 | 0 | 0 | 0 | 0 | 0 |
| Lys | TTT | 81449 | 40102 | 47166 | 27 | 47 | 32 | 3 | 2 | 4 | 0 | 0 | 0 | 0 | 0 | 0 | 0 | 0 | 0 | 0 | 0 | 0 | 0 | 0 | 0 | 0 | 0 | 0 | 0 | 0 | 0 | 0 | 0 | 0 |
| Met | CAT | 122134 | 83729 | 115064 | 176 | 119 | 213 | 9 | 11 | 19 | 2 | 0 | 1 | 0 | 1 | 1 | 0 | 0 | 0 | 0 | 0 | 0 | 0 | 0 | 0 | 0 | 0 | 0 | 0 | 0 | 0 | 0 | 0 | 0 |
| Phe | AAA | 0 | 0 | 0 | 0 | 0 | 0 | 0 | 0 | 0 | 0 | 0 | 0 | 0 | 0 | 0 | 0 | 0 | 0 | 0 | 0 | 0 | 0 | 0 | 0 | 0 | 0 | 0 | 0 | 0 | 0 | 0 | 0 | 0 |
| Phe | GAA | 73674 | 50329 | 56535 | 52 | 42 | 31 | 4 | 5 | 3 | 0 | 0 | 10 | 0 | 0 | 8 | 0 | 0 | 0 | 0 | 0 | 0 | 0 | 0 | 0 | 0 | 0 | 0 | 0 | 0 | 0 | 0 | 0 | 0 |
| Pro | AGG | 64362 | 35241 | 51175 | 131 | 49 | 118 | 24 | 7 | 17 | 2 | 0 | 1 | 2 | 0 | 0 | 0 | 0 | 0 | 0 | 0 | 0 | 0 | 0 | 0 | 0 | 0 | 0 | 0 | 0 | 0 | 0 | 0 | 0 |
| Pro | CGG | 4039 | 1924 | 2166 | 3 | 0 | 1 | 0 | 0 | 0 | 0 | 0 | 0 | 0 | 0 | 0 | 0 | 0 | 0 | 0 | 0 | 0 | 0 | 0 | 0 | 0 | 0 | 0 | 0 | 0 | 0 | 0 | 0 | 0 |
| Pro | TGG | 19765 | 10741 | 15525 | 42 | 33 | 27 | 0 | 2 | 2 | 0 | 0 | 3 | 0 | 0 | 2 | 0 | 0 | 0 | 0 | 0 | 0 | 0 | 0 | 0 | 0 | 0 | 0 | 0 | 0 | 0 | 0 | 0 | 0 |
| Ser | AGA | 228626 | 145814 | 205261 | 239 | 110 | 306 | 47 | 25 | 81 | 6 | 0 | 4 | 0 | 0 | 3 | 0 | 0 | 0 | 0 | 0 | 0 | 0 | 0 | 0 | 0 | 0 | 0 | 0 | 0 | 0 | 0 | 0 | 0 |
| Ser | CGA | 19689 | 17749 | 21887 | 25 | 14 | 39 | 3 | 0 | 0 | 0 | 0 | 0 | 0 | 0 | 0 | 0 | 0 | 0 | 0 | 0 | 0 | 0 | 0 | 0 | 0 | 0 | 0 | 0 | 0 | 0 | 0 | 0 | 0 |
| Ser | GCT | 83449 | 46174 | 61004 | 60 | 24 | 54 | 6 | 2 | 0 | 9 | 0 | 0 | 0 | 0 | 0 | 0 | 0 | 0 | 3 | 0 | 0 | 0 | 0 | 0 | 0 | 0 | 0 | 0 | 0 | 0 | 0 | 0 | 0 |
| Ser | TGA | 74794 | 52775 | 69731 | 48 | 16 | 36 | 9 | 0 | 13 | 0 | 0 | 1 | 3 | 0 | 0 | 0 | 0 | 0 | 0 | 0 | 0 | 0 | 0 | 0 | 0 | 0 | 0 | 0 | 0 | 0 | 0 | 0 | 0 |
| Thr | AGT | 76120 | 43784 | 53703 | 76 | 29 | 121 | 9 | 14 | 13 | 0 | 0 | 0 | 0 | 0 | 0 | 3 | 0 | 0 | 0 | 0 | 0 | 0 | 0 | 0 | 0 | 0 | 0 | 0 | 0 | 0 | 0 | 0 | 0 |
| Thr | CGT | 3019 | 2628 | 2513 | 1 | 2 | 3 | 0 | 0 | 0 | 0 | 0 | 0 | 0 | 0 | 0 | 0 | 0 | 0 | 0 | 0 | 0 | 0 | 0 | 0 | 0 | 0 | 0 | 0 | 0 | 0 | 0 | 0 | 0 |
| Thr | TGT | 226719 | 90263 | 66532 | 99 | 45 | 38 | 4 | 2 | 1 | 0 | 0 | 0 | 0 | 0 | 0 | 0 | 0 | 0 | 0 | 0 | 0 | 0 | 0 | 0 | 0 | 0 | 0 | 0 | 0 | 0 | 0 | 0 | 0 |
| Trp | CCA | 50436 | 29034 | 36424 | 49 | 62 | 35 | 2 | 5 | 1 | 0 | 0 | 0 | 0 | 0 | 0 | 0 | 0 | 0 | 0 | 0 | 0 | 0 | 0 | 0 | 0 | 0 | 0 | 0 | 0 | 0 | 0 | 0 | 0 |
| Tyr | GTA | 11817 | 9834 | 11524 | 27 | 37 | 82 | 20 | 26 | 79 | 31 | 7 | 11 | 0 | 0 | 0 | 0 | 0 | 0 | 0 | 0 | 0 | 0 | 0 | 0 | 0 | 0 | 0 | 0 | 0 | 0 | 0 | 0 | 0 |
| Val | AAC | 43617 | 21142 | 23822 | 32 | 21 | 47 | 4 | 0 | 4 | 0 | 0 | 0 | 0 | 0 | 0 | 0 | 0 | 0 | 0 | 0 | 0 | 0 | 0 | 0 | 0 | 0 | 0 | 0 | 0 | 0 | 0 | 0 | 0 |
| Val | CAC | 6095 | 2944 | 4309 | 35 | 12 | 69 | 0 | 0 | 0 | 0 | 0 | 0 | 0 | 0 | 0 | 0 | 0 | 0 | 0 | 0 | 0 | 0 | 0 | 0 | 0 | 0 | 0 | 0 | 0 | 0 | 0 | 0 | 0 |
| Val | TAC | 47244 | 23873 | 28946 | 35 | 15 | 29 | 4 | 1 | 1 | 0 | 0 | 0 | 0 | 0 | 0 | 0 | 0 | 0 | 0 | 0 | 0 | 0 | 0 | 0 | 0 | 0 | 0 | 0 | 0 | 0 | 0 | 0 | 0 |

| | | premature (U-ending) and mature (CCA-ending) tRNA counts for partial MLP1 knockout (KO) total tRNAs | | | | | | | | | | | | | | | | | | | | | | | | | | | | | | | | |
|---|---|---|---|---|---|---|---|---|---|---|---|---|---|---|---|---|---|---|---|---|---|---|---|---|---|---|---|---|---|---|---|---|---|---|
| **tRNA isotype** | **tRNA anticodon** | **CCA_1** | **CCA_2** | **CCA_3** | **U1_1** | **U1_2** | **U1_3** | **U2_1** | **U2_2** | **U2_3** | **U3_1** | **U3_2** | **U3_3** | **U4_1** | **U4_2** | **U4_3** | **U5_1** | **U5_2** | **U5_3** | **U6_1** | **U6_2** | **U6_3** | **U7_1** | **U7_2** | **U7_3** | **U8_1** | **U8_2** | **U8_3** | **U9_1** | **U9_2** | **U9_3** | **U10_1** | **U10_2** | **U10_3** |
| Ala | AGC | 71772 | 37393 | 39213 | 83 | 32 | 17 | 23 | 17 | 12 | 11 | 2 | 2 | 0 | 0 | 0 | 0 | 0 | 0 | 0 | 0 | 0 | 0 | 0 | 0 | 0 | 0 | 0 | 0 | 0 | 0 | 0 | 0 | 0 |
| Ala | CGC | 21716 | 14584 | 15613 | 11 | 5 | 4 | 0 | 5 | 0 | 0 | 0 | 0 | 0 | 0 | 0 | 0 | 0 | 0 | 0 | 0 | 0 | 0 | 0 | 0 | 0 | 0 | 0 | 0 | 0 | 0 | 0 | 0 | 0 |
| Ala | TGC | 69207 | 31515 | 30878 | 30 | 28 | 23 | 5 | 9 | 0 | 0 | 0 | 0 | 0 | 0 | 0 | 0 | 0 | 0 | 0 | 0 | 0 | 0 | 0 | 0 | 0 | 0 | 0 | 0 | 0 | 0 | 0 | 0 | 0 |
| Arg | ACG | 82876 | 39548 | 53154 | 85 | 45 | 58 | 3 | 0 | 8 | 0 | 0 | 0 | 8 | 16 | 3 | 6 | 0 | 0 | 0 | 0 | 0 | 0 | 0 | 0 | 0 | 0 | 0 | 0 | 0 | 0 | 0 | 0 | 0 |
| Arg | CCG | 1188 | 712 | 692 | 0 | 0 | 0 | 0 | 0 | 0 | 0 | 1 | 0 | 0 | 0 | 0 | 0 | 0 | 0 | 0 | 0 | 0 | 0 | 0 | 0 | 0 | 0 | 0 | 0 | 0 | 0 | 0 | 0 | 0 |
| Arg | CCT | 10167 | 4736 | 5402 | 8 | 5 | 4 | 0 | 0 | 0 | 0 | 0 | 0 | 0 | 0 | 0 | 0 | 0 | 0 | 0 | 0 | 0 | 0 | 0 | 0 | 0 | 0 | 0 | 0 | 0 | 0 | 0 | 0 | 0 |
| Arg | TCG | 4147 | 2343 | 1809 | 4 | 5 | 11 | 0 | 0 | 0 | 0 | 0 | 0 | 2 | 0 | 0 | 0 | 0 | 0 | 0 | 0 | 0 | 0 | 0 | 0 | 0 | 0 | 0 | 0 | 0 | 0 | 0 | 0 | 0 |
| Arg | TCT | 815051 | 508887 | 566999 | 717 | 546 | 491 | 61 | 56 | 30 | 7 | 18 | 51 | 0 | 13 | 5 | 0 | 0 | 0 | 0 | 0 | 0 | 0 | 0 | 0 | 0 | 0 | 0 | 0 | 0 | 0 | 0 | 0 | 0 |
| Asn | GTT | 116604 | 44529 | 53692 | 89 | 34 | 31 | 0 | 0 | 1 | 0 | 0 | 0 | 0 | 0 | 0 | 0 | 0 | 0 | 0 | 0 | 0 | 0 | 0 | 0 | 0 | 0 | 0 | 0 | 0 | 0 | 0 | 0 | 0 |
| Asp | GTC | 120380 | 70875 | 65211 | 93 | 66 | 63 | 23 | 14 | 12 | 38 | 40 | 20 | 6 | 9 | 3 | 0 | 0 | 0 | 0 | 0 | 8 | 0 | 0 | 0 | 0 | 0 | 0 | 0 | 0 | 0 | 0 | 0 | 0 |
| Cys | GCA | 110936 | 75986 | 74105 | 103 | 62 | 84 | 8 | 11 | 7 | 18 | 9 | 6 | 0 | 2 | 0 | 0 | 0 | 1 | 0 | 0 | 0 | 0 | 0 | 0 | 0 | 0 | 0 | 0 | 0 | 0 | 0 | 0 | 0 |
| Gln | CTA | 259803 | 139250 | 158230 | 254 | 129 | 171 | 5 | 1 | 1 | 0 | 40 | 25 | 0 | 0 | 0 | 0 | 0 | 0 | 0 | 0 | 0 | 0 | 0 | 0 | 0 | 0 | 0 | 0 | 0 | 0 | 0 | 0 | 0 |
| Gln | TTA | 99472 | 56703 | 67782 | 425 | 273 | 285 | 14 | 4 | 0 | 32 | 15 | 12 | 0 | 0 | 1 | 0 | 0 | 0 | 0 | 0 | 0 | 0 | 0 | 0 | 0 | 0 | 0 | 0 | 0 | 0 | 0 | 0 | 0 |
| Gln | TTG | 991452 | 578704 | 656541 | 651 | 462 | 447 | 121 | 40 | 57 | 48 | 11 | 49 | 0 | 10 | 16 | 0 | 0 | 0 | 0 | 0 | 0 | 0 | 0 | 0 | 0 | 0 | 0 | 0 | 0 | 0 | 0 | 0 | 0 |
| Glu | CTC | 22690 | 9612 | 8511 | 37 | 30 | 9 | 24 | 11 | 9 | 8 | 2 | 0 | 0 | 0 | 0 | 0 | 0 | 0 | 0 | 0 | 0 | 0 | 0 | 0 | 0 | 0 | 0 | 0 | 0 | 0 | 0 | 0 | 0 |
| Glu | TTC | 144526 | 63140 | 57303 | 202 | 122 | 97 | 18 | 51 | 16 | 0 | 8 | 6 | 0 | 1 | 0 | 0 | 0 | 0 | 0 | 0 | 8 | 0 | 0 | 0 | 0 | 0 | 0 | 0 | 0 | 0 | 0 | 0 | 0 |
| Gly | CCC | 167 | 54 | 86 | 0 | 0 | 0 | 0 | 0 | 0 | 0 | 0 | 0 | 0 | 0 | 0 | 0 | 0 | 0 | 0 | 0 | 0 | 0 | 0 | 0 | 0 | 0 | 0 | 0 | 0 | 0 | 0 | 0 | 0 |
| Gly | GCC | 239527 | 111217 | 108213 | 182 | 47 | 32 | 7 | 9 | 2 | 12 | 6 | 34 | 34 | 15 | 6 | 0 | 0 | 4 | 0 | 0 | 0 | 0 | 0 | 0 | 0 | 0 | 0 | 0 | 0 | 0 | 0 | 0 | 0 |
| Gly | TCC | 31857 | 19319 | 17784 | 35 | 33 | 12 | 11 | 3 | 7 | 0 | 20 | 2 | 10 | 0 | 3 | 0 | 0 | 0 | 0 | 0 | 0 | 0 | 0 | 0 | 0 | 0 | 0 | 0 | 0 | 0 | 0 | 0 | 0 |
| His | GTG | 45057 | 19064 | 21395 | 22 | 26 | 32 | 21 | 5 | 21 | 0 | 23 | 11 | 0 | 0 | 0 | 0 | 0 | 0 | 0 | 0 | 0 | 0 | 0 | 0 | 0 | 0 | 0 | 0 | 0 | 0 | 0 | 0 | 0 |
| Ile | AAT | 328129 | 136498 | 118534 | 224 | 80 | 74 | 23 | 5 | 8 | 0 | 1 | 3 | 0 | 1 | 5 | 9 | 0 | 0 | 0 | 0 | 0 | 0 | 0 | 0 | 0 | 0 | 0 | 0 | 0 | 0 | 0 | 0 | 0 |
| Ile | TAT | 26956 | 12451 | 13939 | 36 | 38 | 21 | 0 | 11 | 8 | 0 | 0 | 1 | 17 | 7 | 5 | 20 | 0 | 0 | 0 | 0 | 0 | 0 | 0 | 0 | 0 | 0 | 2 | 0 | 0 | 0 | 0 | 0 | 0 |
| Leu | AAG | 318831 | 137406 | 155545 | 157 | 85 | 74 | 3 | 11 | 4 | 26 | 5 | 7 | 0 | 4 | 21 | 0 | 0 | 0 | 0 | 0 | 0 | 0 | 0 | 0 | 0 | 0 | 0 | 0 | 0 | 0 | 0 | 0 | 0 |
| Leu | CAA | 1553032 | 485986 | 295666 | 1635 | 732 | 260 | 406 | 220 | 78 | 16 | 26 | 8 | 14 | 6 | 5 | 0 | 9 | 0 | 0 | 0 | 0 | 0 | 0 | 0 | 0 | 0 | 0 | 0 | 0 | 0 | 0 | 0 | 0 |
| Leu | CAG | 300318 | 129195 | 147071 | 124 | 67 | 65 | 3 | 10 | 0 | 12 | 5 | 7 | 0 | 4 | 20 | 0 | 0 | 0 | 0 | 0 | 0 | 0 | 0 | 0 | 0 | 0 | 0 | 0 | 0 | 0 | 0 | 0 | 0 |
| Leu | TAA | 110701 | 56916 | 71315 | 237 | 97 | 123 | 20 | 9 | 7 | 0 | 4 | 6 | 8 | 1 | 4 | 0 | 5 | 3 | 0 | 0 | 0 | 0 | 0 | 0 | 0 | 0 | 0 | 0 | 0 | 0 | 0 | 0 | 0 |
| Leu | TAG | 47973 | 22171 | 24927 | 27 | 12 | 13 | 0 | 3 | 0 | 10 | 10 | 4 | 0 | 3 | 5 | 0 | 0 | 0 | 0 | 0 | 0 | 0 | 0 | 0 | 0 | 0 | 0 | 0 | 0 | 0 | 0 | 0 | 0 |
| Lys | TTT | 107921 | 63792 | 65459 | 27 | 71 | 27 | 17 | 23 | 0 | 13 | 0 | 0 | 14 | 1 | 10 | 0 | 0 | 0 | 0 | 0 | 0 | 0 | 0 | 0 | 0 | 0 | 0 | 0 | 0 | 0 | 0 | 0 | 0 |
| Met | CAT | 124565 | 64897 | 72966 | 110 | 74 | 86 | 10 | 11 | 6 | 0 | 5 | 10 | 2 | 1 | 0 | 0 | 2 | 0 | 0 | 0 | 0 | 0 | 0 | 0 | 0 | 0 | 0 | 0 | 0 | 0 | 0 | 0 | 0 |
| Phe | AAA | 0 | 0 | 0 | 0 | 0 | 0 | 0 | 0 | 0 | 0 | 0 | 0 | 0 | 0 | 0 | 0 | 0 | 0 | 0 | 0 | 0 | 0 | 0 | 0 | 0 | 0 | 0 | 0 | 0 | 0 | 0 | 0 | 0 |
| Phe | GAA | 54969 | 27885 | 31151 | 132 | 97 | 105 | 40 | 28 | 70 | 38 | 22 | 42 | 0 | 4 | 0 | 0 | 0 | 0 | 0 | 4 | 0 | 0 | 0 | 0 | 0 | 0 | 0 | 0 | 0 | 0 | 0 | 0 | 0 |
| Pro | AGG | 27663 | 12788 | 16490 | 102 | 99 | 83 | 18 | 53 | 24 | 4 | 5 | 8 | 8 | 1 | 0 | 0 | 1 | 2 | 0 | 0 | 0 | 0 | 0 | 0 | 0 | 0 | 0 | 0 | 0 | 0 | 0 | 0 | 0 |
| Pro | CGG | 6115 | 2984 | 3118 | 3 | 2 | 2 | 0 | 0 | 0 | 0 | 0 | 0 | 0 | 0 | 0 | 0 | 0 | 0 | 0 | 0 | 0 | 0 | 0 | 0 | 0 | 0 | 0 | 0 | 0 | 0 | 0 | 0 | 0 |
| Pro | TGG | 17471 | 7496 | 8721 | 61 | 17 | 11 | 0 | 13 | 8 | 15 | 71 | 77 | 23 | 10 | 0 | 0 | 0 | 0 | 0 | 0 | 0 | 0 | 0 | 0 | 0 | 0 | 0 | 0 | 0 | 0 | 0 | 0 | 0 |
| Ser | AGA | 224694 | 101158 | 133828 | 157 | 61 | 83 | 22 | 11 | 11 | 22 | 4 | 4 | 0 | 2 | 7 | 0 | 0 | 0 | 0 | 0 | 0 | 0 | 0 | 0 | 0 | 0 | 0 | 0 | 0 | 0 | 0 | 0 | 0 |
| Ser | CGA | 29419 | 14279 | 19091 | 9 | 22 | 10 | 12 | 3 | 1 | 0 | 0 | 0 | 0 | 0 | 0 | 0 | 0 | 3 | 0 | 0 | 0 | 0 | 0 | 0 | 0 | 0 | 0 | 0 | 0 | 0 | 0 | 0 | 0 |
| Ser | GCT | 50080 | 28857 | 25310 | 63 | 28 | 24 | 43 | 9 | 28 | 0 | 0 | 2 | 0 | 1 | 9 | 12 | 1 | 7 | 26 | 0 | 2 | 0 | 0 | 0 | 0 | 0 | 0 | 0 | 0 | 0 | 0 | 0 | 0 |
| Ser | TGA | 111315 | 51610 | 59788 | 52 | 24 | 34 | 21 | 5 | 10 | 33 | 7 | 14 | 0 | 0 | 0 | 0 | 0 | 0 | 0 | 0 | 0 | 0 | 0 | 0 | 0 | 0 | 0 | 0 | 0 | 0 | 0 | 0 | 0 |
| Thr | AGT | 91344 | 56141 | 57326 | 105 | 66 | 81 | 15 | 25 | 11 | 0 | 0 | 0 | 0 | 0 | 7 | 0 | 0 | 2 | 0 | 0 | 0 | 0 | 0 | 0 | 0 | 0 | 0 | 0 | 0 | 0 | 0 | 0 | 0 |
| Thr | CGT | 6923 | 3715 | 3077 | 16 | 5 | 3 | 0 | 0 | 0 | 0 | 0 | 0 | 0 | 0 | 0 | 0 | 0 | 0 | 0 | 0 | 0 | 0 | 0 | 0 | 0 | 0 | 0 | 0 | 0 | 0 | 0 | 0 | 0 |
| Thr | TGT | 369704 | 137505 | 107236 | 168 | 50 | 45 | 4 | 4 | 0 | 0 | 0 | 0 | 0 | 0 | 0 | 0 | 3 | 0 | 0 | 0 | 0 | 0 | 0 | 0 | 0 | 0 | 0 | 0 | 0 | 0 | 0 | 0 | 0 |
| Trp | CCA | 60240 | 33614 | 36027 | 58 | 94 | 64 | 58 | 16 | 18 | 39 | 8 | 9 | 0 | 4 | 5 | 0 | 0 | 0 | 0 | 0 | 0 | 0 | 0 | 0 | 0 | 0 | 0 | 0 | 0 | 0 | 0 | 0 | 0 |
| Tyr | GTA | 16837 | 8815 | 8195 | 112 | 35 | 19 | 96 | 37 | 58 | 172 | 86 | 130 | 20 | 1 | 0 | 0 | 2 | 0 | 0 | 0 | 0 | 0 | 0 | 0 | 0 | 0 | 0 | 0 | 0 | 0 | 0 | 0 | 0 |
| Val | AAC | 42983 | 22066 | 22492 | 263 | 252 | 186 | 83 | 31 | 19 | 7 | 5 | 25 | 0 | 0 | 0 | 0 | 0 | 0 | 0 | 0 | 0 | 0 | 0 | 0 | 0 | 0 | 0 | 0 | 0 | 0 | 0 | 0 | 0 |
| Val | CAC | 3451 | 1898 | 1811 | 105 | 103 | 62 | 0 | 2 | 0 | 11 | 2 | 0 | 0 | 0 | 0 | 0 | 0 | 0 | 0 | 0 | 0 | 0 | 0 | 0 | 0 | 0 | 0 | 0 | 0 | 0 | 0 | 0 | 0 |
| Val | TAC | 24016 | 12533 | 12597 | 71 | 80 | 100 | 3 | 3 | 15 | 0 | 0 | 0 | 0 | 0 | 0 | 9 | 0 | 0 | 0 | 0 | 0 | 0 | 0 | 0 | 0 | 0 | 0 | 0 | 0 | 0 | 0 | 0 | 0 |

**Table S4.** Number of tRNA genes encoded in the genome of different eukaryotic species for each isotype (top) and isoacceptor (bottom). The number of tRNA genes was obtained by counting the number of entries for each tRNA isotype and anticodon from the Genomic tRNA Database (GtRNAdb) and from the UCSC Genome Browser for *Tetrahymena thermophila* in Additional Table 3. NNN anticodons were excluded from the analysis.

| tRNA isotype | *T. thermophila* | *S. pombe* | *S. cerevisiae* | *A. thaliana* | *D. melanogaster* | *M. musculus* | *H. sapiens* |
|---|---|---|---|---|---|---|---|
| **Ala** | 44 | 12 | 16 | 33 | 17 | 46 | 45 |
| **Arg** | 36 | 13 | 19 | 37 | 26 | 25 | 31 |
| **Asn** | 31 | 6 | 10 | 16 | 10 | 7 | 36 |
| **Asp** | 31 | 8 | 16 | 27 | 14 | 22 | 20 |
| **Cys** | 16 | 3 | 4 | 16 | 7 | 62 | 32 |
| **Gln** | 52 | 6 | 10 | 17 | 12 | 19 | 31 |
| **Glu** | 44 | 10 | 16 | 27 | 21 | 22 | 28 |
| **Gly** | 42 | 12 | 21 | 41 | 20 | 32 | 36 |
| **His** | 17 | 4 | 7 | 10 | 6 | 11 | 10 |
| **Ile** | 41 | 9 | 15 | 22 | 12 | 18 | 24 |
| **iMet** | 0 | 4 | 5 | 2 | 6 | 10 | 10 |
| **Leu** | 55 | 13 | 21 | 42 | 22 | 29 | 38 |
| **Lys** | 29 | 12 | 21 | 32 | 19 | 41 | 40 |
| **Met** | 33 | 3 | 5 | 23 | 6 | 9 | 10 |
| **Phe** | 38 | 5 | 10 | 16 | 8 | 7 | 17 |
| **Pro** | 28 | 9 | 12 | 67 | 17 | 20 | 25 |
| **Ser** | 46 | 13 | 17 | 64 | 20 | 22 | 28 |
| **Sup** | 0 | 0 | 0 | 1 | 0 | 0 | 1 |
| **Thr** | 29 | 10 | 16 | 23 | 18 | 1 | 22 |
| **Trp** | 21 | 3 | 6 | 14 | 8 | 0 | 7 |
| **Tyr** | 19 | 4 | 8 | 76 | 10 | 0 | 16 |
| **Val** | 32 | 12 | 18 | 30 | 15 | 0 | 34 |
| **Sum** | **684** | **171** | **273** | **636** | **294** | **403** | **541** |

| tRNA isotype | anticodon | *T. thermophila* | *S. pombe* | *S. cerevisiae* | *A. thaliana* | *D. melanogaster* | *M. musculus* | *H. sapiens* |
|---|---|---|---|---|---|---|---|---|
| **Ala** | | **44** | **12** | **16** | **33** | **17** | **46** | **45** |
| | **AGC** | 31 | 9 | 11 | 16 | 12 | 22 | 30 |
| | **CGC** | 2 | 1 | | 7 | 3 | 10 | 5 |
| | **GGC** | | | | | | 3 | |
| | **TGC** | 11 | 2 | 5 | 10 | 2 | 11 | 10 |
| **Arg** | | **36** | **13** | **19** | **37** | **26** | **25** | **31** |
| | **ACG** | 8 | 8 | 6 | 9 | 10 | 6 | 7 |
| | **CCG** | 1 | 1 | 1 | 4 | | 3 | 4 |
| | **CCT** | 2 | 1 | 1 | 8 | 3 | 5 | 8 |
| | **TCG** | 1 | 1 | | 6 | 10 | 5 | 6 |
| | **TCT** | 24 | 2 | 11 | 10 | 3 | 6 | 6 |
| **Asn** | | **31** | **6** | **10** | **16** | **10** | **7** | **36** |
| | **ATT** | | | | | | | 2 |
| | **GTT** | 31 | 6 | 10 | 16 | 10 | 13 | 35 |
| **Asp** | | **31** | **8** | **16** | **27** | **14** | **22** | **20** |
| | **GTC** | 31 | 8 | 16 | 27 | 14 | 16 | 19 |
| **Cys** | | **16** | **3** | **4** | **16** | **7** | **62** | **32** |
| | **ACA** | | | | | | 1 | 1 |
| | **GCA** | 16 | 3 | 4 | 16 | 7 | 61 | 31 |
| **Gln** | | **52** | **6** | **10** | **17** | **12** | **19** | **31** |
| | **CTA *** | 9 | | | | | | |
| | **CTG** | | 2 | 1 | 9 | 8 | 12 | 22 |
| | **TTA *** | 30 | | | | | | |
| | **TTG** | 13 | 4 | 9 | 8 | 4 | 7 | 9 |
| **Glu** | | **44** | **10** | **16** | **27** | **21** | **22** | **28** |
| | **CTC** | 7 | 6 | 2 | 13 | 15 | 14 | 12 |
| | **TTC** | 37 | 4 | 14 | 14 | 6 | 8 | 16 |
| **Gly** | | **42** | **12** | **21** | **41** | **20** | **32** | **36** |
| | **ACC** | | | | | | 2 | |
| | **CCC** | 1 | 1 | 2 | 5 | | 7 | 10 |
| | **GCC** | 29 | 8 | 16 | 23 | 14 | 15 | 15 |
| | **TCC** | 12 | 3 | 3 | 13 | 6 | 8 | 11 |
| **His** | | **17** | **4** | **7** | **10** | **6** | **11** | **10** |
| | **ATG** | | | | | | 1 | |
| | **GTG** | 17 | 4 | 7 | 10 | 6 | 10 | 10 |
| **Ile** | | **41** | **9** | **15** | **22** | **12** | **18** | **24** |
| | **AAT** | 29 | 8 | 13 | 17 | 10 | 12 | 16 |
| | **GAT** | | | | | | 1 | 3 |
| | **TAT** | 12 | 1 | 2 | 5 | 2 | 5 | 5 |
| **iMet** | | | **4** | **5** | **2** | **6** | **10** | **10** |
| | **CAT** | | 4 | 5 | 2 | 6 | 10 | 10 |
| **Leu** | | **55** | **13** | **21** | **42** | **22** | **29** | **38** |
| | **AAG** | 19 | 5 | | 12 | 4 | 5 | 13 |
| | **CAA** | 11 | 4 | 10 | 10 | 4 | 4 | 7 |
| | **CAG** | 2 | 1 | | 3 | 8 | 10 | 9 |
| | **GAG** | | | 1 | 1 | | | |
| | **TAA** | 19 | 2 | 7 | 6 | 4 | 7 | 5 |
| | **TAG** | 4 | 1 | 3 | 10 | 2 | 3 | 4 |
| **Lys** | | **29** | **12** | **21** | **32** | **19** | **41** | **40** |
| | **CTT** | | 9 | 14 | 18 | 13 | 26 | 21 |
| | **TTT** | 29 | 3 | 7 | 14 | 6 | 15 | 19 |
| **Met** | | **33** | **3** | **5** | **23** | **6** | **9** | **10** |
| | **CAT** | 33 | 3 | 5 | 23 | 6 | 9 | 10 |
| **Phe** | | **38** | **5** | **10** | **16** | **8** | **7** | **17** |
| | **AAA** | 1 | | | | | | |
| | **GAA** | 37 | 5 | 10 | 16 | 8 | 7 | 17 |
| **Pro** | | **28** | **9** | **12** | **67** | **17** | **20** | **25** |
| | **AGG** | 20 | 6 | 2 | 16 | 7 | 8 | 11 |
| | **CGG** | 1 | 1 | | 5 | 5 | 3 | 4 |
| | **GGG** | | | | | | 1 | 1 |
| | **TGG** | 7 | 2 | 10 | 45 | 5 | 8 | 9 |
| **Ser** | | **46** | **13** | **17** | **64** | **20** | **22** | **28** |
| | **ACT** | | | | | | | 1 |
| | **AGA** | 20 | 7 | 11 | 37 | 8 | 9 | 11 |
| | **CGA** | 2 | 1 | 1 | 4 | 4 | 3 | 4 |
| | **GCT** | 15 | 3 | 2 | 13 | 6 | 9 | 8 |
| | **GGA** | | | | 1 | | | |
| | **TGA** | 9 | 2 | 3 | 9 | 2 | 1 | 4 |
| **Sup** | | | | | **1** | | | **1** |
| | **TTA** | | | | 1 | | | 1 |
| **Thr** | | **29** | **10** | **16** | **23** | **18** | **1** | **22** |
| | **AGT** | 22 | 7 | 11 | 10 | 9 | 1 | 10 |
| | **CGT** | 1 | 1 | 1 | 5 | 3 | | 6 |
| | **TGT** | 6 | 2 | 4 | 8 | 6 | | 6 |
| **Trp** | | **21** | **3** | **6** | **14** | **8** | | **7** |
| | **CCA** | 21 | 3 | 6 | 14 | 8 | | 7 |
| **Tyr** | | **19** | **4** | **8** | **76** | **10** | | **16** |
| | **ATA** | | | | | | | 1 |
| | **GTA** | 19 | 4 | 8 | 76 | 10 | | 15 |
| **Val** | | **32** | **12** | **18** | **30** | **15** | | **34** |
| | **AAC** | 21 | 9 | 14 | 15 | 6 | | 11 |
| | **CAC** | 5 | 1 | 2 | 8 | 7 | | 18 |
| | **TAC** | 6 | 2 | 2 | 7 | 2 | | 5 |
| **Sum** | | **684** | **171** | **273** | **636** | **294** | **403** | **541** |

* Anticodons encoding stop codons in eukaryotes except in *T. thermophila*

**Table S5.** List of oligonucleotides used in this study.

| **gBlock Gene Fragments used as DNA template for cloning MLP1** | |
|---|---|
| gBlock MLP1 CDS codon optimized | ATGTCAATCAATAAAGAAGAGGTAAAAAAGCAAGTCGAATACTATCTGTCGGACAAGAACTTGGTAAATGATGAGAAATTTCGCACTATCATTCAAGAGCACCCAGAG<br>GGGTATTTATCCTTCGGCAATATCTTAAACTGTAATAAGATTAAGCGTCTGGGTGTGACGACATTCGAACAACTGGCAACGTCCCTTGCTGATAGTACCTTAGTTGAA<br>TTAAATGAGGCAAAGGACTCTGTGCGTCGTGCCGGAAATAAACCCATTCCAGCGAAGGAAGCAGTTGACCCCGCTGAAGCTGCAGAAAAAGAAGCGCGTGAGGC<br>CGAAAAAAAGGAACTGATCAACTTTTATGAAACTTTTCAACCGATTATCTTCTCAACCGCCTGCGAACAAGAGGGGGTAGCGAACTGGCGCAACATCACAGAGGCCT<br>TGCTGAAGCAACATAACGTTCATGCTCCGTATTGCCGCTTTGGCAAGTTGGAGGGTAACTTTGCATTGAATAAAGACAAAACTTCGCAAGAGGTTATCGACCAGCTT<br>GTGCAAGACGGGCTTCAATTTGGAGAGTCTAAGGTGACTATCAAAGTGAGTGAGGGCGAAGCCCTGTCAAAGTTTTGGGAGCTGCATGGACGCCACTACAACGGT<br>GTGATGGAATTGAAAAAGAAGGAGGTCAATCAGACAGTTAAAGCTAAGAAAGATAAAAAGGAGAAAAAACAAAAACGTGAATTCGAATTCGGGGGAGAAAAATATAC<br>AGATGTATTGACTATTAAAAATTTGTTTAAGGGAATCTTAGGGCGTACGGCGAATGGACAAAAGATCATTAGTCCTTATCACGAGATGTTGAAGAGCCTTCTTGAGTA<br>TCACAATAACAAAGAAGCCAAATTAAAAGATTTAGATCACTTCACGGTAGACGTCCATCCTGAGCATAAAGATACACGTTGTTTCTTTGTCGTCAAATCGGACGGAAC<br>GAAGGAAGATTTCTCAGCGGTTAAGTGCATCTCAAACTTTGAGGAAAAACTTAAACTTTGA |

| **Oligonucleotides used for cloning MLP1 and MLP1-mutants in pET28a (bacterial expression - protein purification) and pRep4 (yeast expression - tRNA mediated suppression assay).** Restriction enzyme cleavage sites are underlined. 6X His tag is shown in bold and blue. Point mutations are shown in bold and pink. | | | |
|---|---|---|---|
| MLP1 SalI 6X His For | pRep4 | 5' | GCGCGGTCGACATG**CACCATCACCATCACCAT**TCAATCAATAAAGAAGAGGTAAAAAAGC |
| MLP1 Q11A/Y14A SalI 6X His For | pRep4 | 5' | GCGCGGTCGACATG**CACCATCACCATCACCAT**TCAATCAATAAAGAAGAGGTAAAAAAG**GCA**GTCGAA**GCC**TATCTGTCGGACAAGAACTTGGTAAATG |
| MLP1 95 SalI 6X His For | pRep4 | 5' | GCGCGGTCGACATG**CACCATCACCATCACCAT**GACCCCGCTGAAGCTGCAGAAAAAGAAGCGCGTG |
| MLP1 NheI For | pET28a | 5' | GCGCGGCTAGCATGTCAATCAATAAAGAAGAGGTAAAAAAGCAAGTC |
| MLP1 Q11A/Y14A NheI For | pET28a | 5' | GCGCGGCTAGCATGTCAATCAATAAAGAAGAGGTAAAAAAG**GCA**GTCGAA**GCC**TATCTGTCGGACAAGAACTTGG |
| MLP1 95 NheI For | pET28a | 5' | GCGCGGCTAGCATGGACCCCGCTGAAGCTGCAGAAAAAGAAGCGCGTG |
| MLP1 BamHI Rev | pET28a/pRep4 | 5' | GCGCGGGATCCTCAAAGTTTAAGTTTTTCCTCAAAGTTTGAG |
| MLP1 95 BamHI Rev | pET28a/pRep4 | 5' | GCGCGGGATCCTCAGTCAACTGCTTCCTTCGCTGGAATGGGTTTATTTCCG |
| MLP1 250 BamHI Rev | pET28a/pRep4 | 5' | GCGCGGGATCCTCAATCTGTATATTTTTCTCCCCCGAATTCGAATTC |

| **Oligonucleotides used to generate T7 DNA template for *in vitro* transcription of tRNAs for electromobility shift assays (EMSAs).** T7 promotor sequence is shown in bold and blue. | | |
|---|---|---|
| Leu AAG DNA template | 5' | GATGAAGTGGCCGAGCGGTCTAAGGCGGTAGATTAAGGCTCTATTCCGAAAGGGCGCGAGTTCGAATCTCGCCTTCATCA |
| Leu AAG **T7 promoter** + 5'-leader For | 5' | **GCTAATACGACTCACTATA**gagcgaaGATGAAGTGGCCGAGCGGTC |
| Leu AAG **T7 promoter** + mature For | 5' | **GCTAATACGACTCACTATA**GATGAAGTGGCCGAGCGGTC |
| Leu AAG 3'-trailer + $(U)_5$ Rev | 5' | aaaaatgTGATGAAGGCGAGATTCGAAC |
| Leu AAG mature Rev | 5' | TGATGAAGGCGAGATTCGAAC |
| Gln TTA DNA template | 5' | GGTTCCATAGTATAGTGGTTAGTACTGGGGACTTTAAATCCCTTGACCTGGGTTCGAATCCCAGTGGGACCT |
| Gln TTA **T7 promoter** + 5'-leader For | 5' | **GCTAATACGACTCACTATA**gcatgtccGGTTCCATAGTATAGTGGTTAGTAC |
| Gln TTA **T7 promoter** + mature For | 5' | **GCTAATACGACTCACTATA**GGTTCCATAGTATAGTGGTTAGTAC |
| Gln TTA 3'-trailer + $(U)_5$ Rev | 5' | aaaaagcAGGTCCCACTGGGATTCGAACCC |
| Gln TTA mature Rev | 5' | AGGTCCCACTGGGATTCGAACCC |

| **Oligonucleotides used for cloning MLP1 partial knockout strain** | | | |
|---|---|---|---|
| MLP1 5'-flanking KpnI For | pNeo4 | 5' | CGCGGGTACCGGTATGTTGTTTGAGATCTTATTAGAATCAC |
| MLP1 5'-flanking XhoI Rev | pNeo4 | 5' | CGCGCTCGAGTTATTCAAAAGTAGTAACTCCGAGTCTCTT |
| MLP1 3'-flanking NotI For | pNeo4 | 5' | CGCGGCGGCCGCTCACGAAATGGTAAATAAATAAATACGATT |
| MLP1 3'-flanking SacI Rev | pNeo4 | 5' | ATTAGAGCTCAGTATATTAAAAGCAATCAATAAATATAAA |

| **DNA probes used for northern blotting.** Anticodon sequence shown in bold and 5'-leader, 3'-trailer and intron sequences in lowercase if applicable. | | |
|---|---|---|
| *T. thermophila* U5 snRNA | 5' | CACATTGAAAAACCCAGCTCACGG |
| *T. thermophila* 5.8S rRNA | 5' | CTGCAATTCGCATTGCGT |
| *T. thermophila* Tyr GTA pre-tRNA 3'-trailer | 5' | aaaaaaaaTCCGAACTCCCGGG |
| *T. thermophila* Tyr GTA mature tRNA | 5' | GACCACCGGATTTACAGTCCG |
| *T. thermophila* Ile TAT pre-tRNA intron | 5' | GACacacttttataaggttggtcagcaaaT**ATA**AATC |
| *T. thermophila* Ile TAT/Leu TAA pre-tRNA 3'-trailer | 5' | aaaaaaaaagTAGCTCAGAGAGGGTTC |
| *T. thermophila* Ile TAT/Leu TAA pre-tRNA 5'-leader | 5' | ACCACTGAGCCtttcaaag |
| *T. thermophila* Leu TAA pre-tRNA intron | 5' | GACactcttttctaaatgtaggaggatgaacG |
| *T. thermophila* Val CAC pre-tRNA intron | 5' | TTAAGGTTtattagaccttC**GTG**ATA |
| *T. thermophila* Val CAC pre-tRNA 3'-trailer | 5' | aaaaaaGTCGACACTAGGGTTTGAA |
| *T. thermophila* Val CAC pre-tRNA 5'-leader | 5' | ACTACTTCACCATGCCGACtatagcag |
| *T. thermophila* Arg TCT mature tRNA | 5' | CAACCAAGGTGGGACTCGAACCCAC |
| *S. pombe* U5 snRNA | 5' | CTGGTAAAAGGCAAGAAACAGATACG |
| *S. pombe* suppressor Ser TCA pre-tRNA intron | 5' | gaatacaggaT**TGA**AGTCT |
| *S. pombe* suppressor Ser TCA pre-tRNA intron (unlabelled probe) | 5' | gaatacaggaT**TCA**AGTCT |
| *S. pombe* suppressor Ser TCA mature tRNA | 5' | **TGA**AGTCTAACTCCTT |
| *S. pombe* suppressor Ser TCA mature tRNA (unlabelled probe) | 5' | **TCA**AGTCTAACTCCTT |
| *S. pombe* Lys CTT pre-tRNA intron | 5' | CTTCTGATaccattcgT**AAG**AGTC |
| *S. pombe* Lys CTT pre-tRNA 3'-trailer | 5' | aaattaaccTCCCAAG |
| *S. pombe* Lys CTT pre-tRNA 5'-leader | 5' | ATTGAGCCACTCGGGAcgcgtt |
| *S. pombe* Lys CTT mature tRNA | 5' | CTCCCAAGGCGAGACTCGAACTCGCAA |

| **DNA probes used for pre-tRNA intron-specific PCR amplification for Sanger sequencing RIP-3'-RACE** | | |
|---|---|---|
| Lys CTT pre-tRNA intron | 5' | TCTGACTCTTATGATGGTAATCAGA |
| Tyr GTA pre-tRNA intron | 5' | CCGGCTGTAATTATAAAGATACC |

| **RNA for electromobility shift assays (EMSAs)** | | |
|---|---|---|
| U10 | 5' | rUrUrUrUrUrUrUrUrUrU |
| U10-3'-P | 5' | rUrUrUrUrUrUrUrUrUrU/3Phos/ |
| 4U (wild type) | 5' | rCrUrGrCrUrGrUrUrUrU |
| $U_1C$ | 5' | rCrUrGrCrUrGrUrUrUrC |
| $U_2C$ | 5' | rCrUrGrCrUrGrUrUrCrU |
| $U_3C$ | 5' | rCrUrGrCrUrGrUrCrUrU |
| 4C | 5' | rCrUrGrCrIrGrCrCrCrC |

**Table S6**. NCBI accession numbers of primary amino acid sequences used for conservation analysis

| | | La | Rex1p | RNase Z / ELAC2 | LSm2 | LSm3 | LSm4 | LSm5 | LSm6 | LSm7 | LSm8 |
|---|---|---|---|---|---|---|---|---|---|---|---|
| **Vertebrates** | *Homo sapiens* | NP_003133.1 | NP_065746.3 | NP_060597.4 | NP_067000.1 | NP_055278.1 | NP_036453.1 | NP_036454.1 | NP_009011.1 | NP_057283.1 | NP_057284.1 |
| | *Mus musculus* | NP_033304.1 | NP_080128.2 | NP_001349912.1 | NP_001103571.1 | NP_080585.1 | NP_056631.2 | NP_079796.1 | NP_001177933.1 | NP_001346078.1 | NP_598700.1 |
| | *Xenopus laevis* | NP_001081021.1 | | | | | | | | | |
| | *Danio rerio* | NP_955841 | NP_001119888.1 | NP_001243133.1 | | | | | | | |
| **Invertebrates** | *Caenorhabditis elegans* | NP_491411.1 | NP_498135.2 | NP_001023110.1 | | | | | | | |
| | *Drosophila melanogaster* | NP_477014.1 | NP_001034073.1 | NP_724916.1 | | | | | | | |
| **Amoebozoa** | *Dictyostelium discoideum* | XP_640625.1 | | | | | | | | | |
| **Kinetoplastide** | *Trypanosoma brucei* | XP_822491.1 | | | | | | | | | |
| **Embryophytes** | *Arabidopsis thaliana* | NP_567904.1 | | | | | | | | | |
| | | NP_178106.2 | | | | | | | | | |
| | *Physcomitrella patens* | XP_024359573.1 | | | | | | | | | |
| **Chlorophytes** | *Chlamydomonas reinhardtii* | XP_042914771.1 | | | | | | | | | |
| **Fungi** | *Schizosaccharomyces pombe* | NP_593315.1 | NP_594627.2 | NP_595514.1 | NP_588459.1 | NP_595747.1 | NP_001342832.1 | NP_596373.1 | NP_594380.1 | NP_588340.1 | NP_588509.1 |
| | *Saccharomyces cerevisiae* | NP_010232.1 | | | NP_009527.1 | NP_013543.3 | NP_011037.3 | NP_011073.1 | NP_010666.2 | NP_014252.2 | NP_012556.2 |
| **Alveolates** | *Paramecium tetraurelia* | XP_001439258.1 | XP_001435945.1 | XP_001346883.1 | | | | | | | |
| | *Tetrahymena thermophila* | XP_001019287.2 | XP_001033374.2 | XP_001031902.2 | XP_012654399 | XP_012656392.1 | XP_001008519.2 | XP_012655775.1 | XP_012653392.1 | XP_001031297.1 | XP_001470817.1 |
